# Editing-independent effects of Adar in *Drosophila melanogaster*

**DOI:** 10.1101/2025.09.13.675992

**Authors:** Khadija Hajji, Damiano Amoruso, Barbora Nováková, Nagraj Sambrani, Deying Yang, Anzer Khan, Vojtech Bystry, Alejandro Medaglia-Mata, Domenico Alessandro Silvestris, Ernesto Picardi, Mary A. O’Connell, Liam P. Keegan

## Abstract

Vertebrate ADAR RNA editing enzymes prevent cellular dsRNA from aberrantly activating antiviral dsRNA sensors. ADARs inhibit antiviral sensor activation by deaminating selected adenosines to inosines in dsRNA and by engaging in inhibitory protein interactions with sensors on dsRNA. ADARs interact with Dicers and, in the *Drosophila Adar^5G1^* null mutant Dicer 2 acts as the antiviral dsRNA sensor mediating aberrant innate immune induction. We overexpressed active Adar isoforms or a catalytically-inactive Adar E374A protein from *UAS-Adar* constructs under the control of a temperature-regulated *Act5C^ts^-GAL4* driver. Overexpression of the edited AdarG isoform or AdarEA cause larval lethality with aberrant innate immune induction. Some escaper pupae are formed with head eversion defects. AdarS overexpression is also lethal with no progeny pupae. Ecdysone signaling gene and innate immune gene expression are aberrantly elevated in the AdarG overexpressing pupae. RNAi knockdown of *Ecdysone Receptor A* (*EcR A*) or increasing expression of HP1 partially rescue AdarG overexpression defects and normalize gene expression in progeny flies, indicating that aberrant epigenetic silencing is also involved. The structure of an ADAR2 dimer on dsRNA shows the the glycine in AdarG is suitably positioned for Adar contacts with other proteins on dsRNA.

## Introduction

ADAR RNA editing enzymes deaminate specific adenosines to inosines within double-stranded (ds)RNA. Editing of cellular dsRNA prevents aberrant activation of antiviral dsRNA sensors and aberrant interferon induction (Mannion et al. 2014; Liddicoat et al. 2015). Human *ADAR* mutants cause Aicardi-Goutières Syndrome 6, a virus infection mimic syndrome caused by aberrant interferon induction that can lead to fatal encephalitis (Rice et al. 2012). Vertebrate ADAR1 protein further inhibits aberrant activation of antiviral dsRNA sensors by direct ADAR1-sensor protein interactions on the dsRNA (Sinigaglia et al. 2024). Such editing-independent ADAR1 interactions with the dsRNA sensor Protein Kinase R (PKR) (Sinigaglia et al. 2024) and with the antiviral Z-RNA sensor Z-RNA/DNA-binding protein 1 (ZBP1) (Hubbard et al. 2022) are important to prevent aberrant sensor activation. It is likely that editing-independent effects of ADAR proteins may be observed also in *Drosophila* where they can be dissected genetically.

*Drosophila melanogaster* has a single adenosine deaminase acting on RNA (*Adar*) gene on the distal X-chromosome (Palladino et al. 2000a). The site-specific recoding type of Adar editing is more widespread in *Drosophila* compared to mammals (Graveley et al. 2011). However, the *Adar^5G1^*null mutant also shows aberrant innate immune gene induction (Deng et al. 2020), in addition to other defects likely related to loss of CNS recoding editing, such as partial embryonic and larval lethality, locomotion defects in larvae and flies (Palladino et al. 2000b), aberrant presynaptic vesicle accumulation in neurons and increased neuronal excitibilty (Maldonado et al. 2013), aberrant autophagy and age-dependent vacuolar brain neurodegeneration (Khan et al. 2020).

*Adar* transcript expression is much lower in embryos and larvae and increases significantly at metamorphosis (Palladino et al. 2000b; Graveley et al. 2011), to edit approximately 1300 editing sites in over six hundred ORFs, especially in the brain which grows and becomes much more complex during metamorphosis. Especially from metamorphosis onwards, Adar also edits the *Adar S/G* editing site, where a serine codon is edited to a glycine codon to generate AdarS and AdarG isoforms expressed at approximately a 60:40 ratio in flies (Palladino et al. 2000a). The edited AdarG enzyme isoform edits approximately eightfold less efficiently than the genome-encoded AdarS isoform *in vitro* at 37℃ (Keegan et al. 2005). In flies at 25℃ AdarG also edits somewhat more weakly than AdarS when isoforms are expressed from *UAS-cDNA* constructs under the control of GAL4 drivers (Keegan et al. 2005). Across a large number of editing sites, AdarG edited somewhat less that AdarS when the endogenous *Adar* gene was modified to express only one of the two isoforms (Savva et al. 2012); however, AdarG also edited some sites better than AdarS (Savva et al. 2012). Consistent with the higher editing activity of AdarS, overexpression of this isoform from randomly inserted *UAST-Adar^S^* cDNA constructs under the temperature-dependent control of the strongly and ubiquitously expressed *Act5C^ts^-GAL4* driver from the embryonic stage onwards was embryonic and larval lethal, whereas overexpression of AdarG or a catalytically inactive Adar E374A protein were not (Keegan et al. 2005). We obtained editing-dependent embryonic and larval lethality by overexpressing AdarS under temperature-dependent *Act5C^ts^-GAL4* driver control and obtained suppressors that reduced the aberrantly decreased neuronal excitability of *Act5C^ts^> Adar^S^* (Li et al. 2014).

Aberrant antiviral innate immune induction occurs in *Adar^5G1^*mutant brains and is prevented by RNAi knockdown of Dicer-2 (Deng et al. 2020), indicating that Dcr2 acts as the innate immune sensor for aberrantly unedited cellular dsRNA in *Adar^5G1^*, as it does during dsRNA virus infection (Deddouche et al. 2008). ADARs, expecially the heterologously expressed, cytoplasmic human ADAR1 p150, also antagonize Dcr2-mediated cytoplasmic RNAi in an editing-independent manner in *Drosophila* (Heale et al. 2009). AdarS assists and AdarG reduces HP1 expression and heterochromatin silencing of transposons (Savva et al. 2013); this type of heterochromatin silencing also involes dsRNA formation at the chromosomal locus to be targeted, with subsequent involvement of and Dcr2 and Ago2.

The lethality of the *Adar^5G1^* null mutant is incomplete and variable and is difficult to use for genetic screens for suppressors of *Adar* mutant lethality (Khan et al. 2020). To generate mutant phenotypes more suitable for genetic screens for *Adar-*modifier mutants we again experimentally increased Adar expression. To elucidate the role of the AdarG isoform in particular, we constructed new *UASg-Adar^G^*plasmids, inserted these at phiC30 *attP2* and *attP40* landing sites and again overexpressed AdarG at higher levels under *Act5C-GAL4* driver control (Keegan et al. 2005). This overexpression from *attP2 UASg-Adar^G^* leads to lethality at embryonic and larval stages, and placing this under GAL80^ts^ control gave similar lethality and also a pupal ecdysone-related defect and failure of eclosion that were not observed previously with Adar isoforms expressed at lower levels. The AdarG overexpressing pupae have aberrant expression of transcripts encoding ecdysone regulatory proteins and innate immune antimicrobial peptides (AMPs). Significantly, all AdarG overexpression phenotypes are mimicked by similar overexpression of catalytically inactive AdarE367A protein, whereas similar AdarS overexpression gives pre-pupal lethality but no pupae. The AdarEA overexpression lethality indicates that AdarG overexpression lethality does not require Adar RNA editing activity and is likely to involve Adar interactions with other proteins. A genetic screen for suppressors of this AdarG overexpression lethality revealed rescue by RNAi knockdown of the *EcRA* transcript encoding Ecdysone Receptor isoform A. Adar is normally expressed in the prothoracic gland (PG) in the ring gland, where larval ecdysone is synthesized; however, *phantom22 (phm22)*-*GAL4*-driven AdarG overexpression in PG cells leads to giant larvae that pupate late or never. Transcripts encoding ecdysone-related enzymes and signaling proteins are extremely reduced in these larvae, including transcripts of the centric heterochromatin *spookier* and *neverland* genes that require the heterochromatin protein HP1 for full expression (Ohhara et al. 2022). Increased expression of HP1 also partially rescues *AdarG* overexpression lethality in the flies overexpressing AdarG ubiquitously. Therefore Adar RNA editing of its own transcript results in the expression of AdarS and AdarG isoforms differing in one amino acid that exhibit very different effects when they are overexpressed. The serine or glycine residue is predicted to be suitably located to engage in Adar interactions with other proteins on dsRNA.

## Results

### AdarG isoform overexpression throughout development causes embryonic and larval lethality

We sought to have stronger expression of AdarG and other isoforms with *UAS-Adar* cDNA constructs having an improved Kozak consensus sequence for translation and more consistent expression levels for different Adar isoforms by inserting all constructs at the same defined phiC30 recombinase *attP* landing sites on chromosomes II and III. New transgenic lines were generated in which an *UAS−Adar^G^* construct or an *UAS−Adar^E367A^* construct encoding a deaminase-inactive protein lacking RNA editing activity were inserted, initially specifically at the *attP2* landing site on chromosome III by phiC30-mediated recombination. Subsequently a complete series of chromosome II *attP40* lines were generated for expression of *Drosophila* Adar isoforms and mutant proteins.

*Act5C−GAL4*-driven overexpression of the edited adult AdarG isoform expressed from the *attP2 UASG-Adar^G^*construct insertion in Chromosome III or from a similar *attP40* insertion on chromosome II resulted in complete lethality before the pupal stage. To determine whether the lethality observed in AdarG overexpressing pupae is caused by an editing-dependent or editing-independent effect of Adar, the catalytically inactive Adar E374A (hereafter AdarEA) mutant (Deng et al. 2020) was overexpressed in *Act5C> attP40-Adar^EA^* progeny. Similar to what was observed with AdarG, overexpression of AdarEA resulted in pre-pupal lethality, indicating that the effect is due to Adar protein expression and does not require Adar RNA editing activity.

### RNAi knockdown screen for suppressors of AdarG-overexpression pre-pupal lethality and of a new pupal head eversion defect

To enable genetic screening for suppressors of the *Act5C> Adar^G^* lethal phenotype, a temperature-sensitive *GAL4-GAL80^ts^* system was employed. A stable strain was generated in which the Chromosome II *Act5C−GAL4* construct insertion and a *Tubulin(Tub)−GAL80^ts^* construct insertion were combined (*Act5C^ts/^/CyO* strain). The *Act5C^ts^* recombinant second chromosome was further combined with *attP2 UAS−Adar^G^*on Chr III to make the *w^1118^; Act5C−GAL4, Tub-GAL80^ts^(T7) /SM5, CyO; attP2 UASg−Adar^G^*(hereafter called *Act5C^ts^/CyO> Adar^G^*) strain. The GAL80^ts^ protein inhibits GAL4 activity in a temperature-dependent manner (McGuire et al. 2004), allowing the *Act5C^ts^/CyO> Adar^G^* tester strain for the suppressor screen to be propagated at 18 °C. When flies of this strain obtained at 18 °C are mated to each other and moved to 25 °C the parents survive and lay eggs but all the progeny overexpressing AdarG die as embryos or larvae.

When *Act5C^ts^/CyO> Adar^G^* virgin females are crossed with *w^1118^* wild-type or *UAS-GFP* flies at 25°C the resulting non-*Curly*, *Act5C^ts^> Adar^G^* progeny carrying both *Act5C^ts^-GAL4* and *UAS-Adar^G^* (AdarG overexpressing) mostly die before the pupal stage but also generate some pupae, probably because the GAL80^ts^ protein still partially inhibits transcription activation by GAL4 at 25^°^C. All of these escaper pupae show a classic pupal head eversion defect, including truncated legs, reduced wing size, extended abdomens, and failure of head eversion, with the head remaining embedded within the thorax (Thummel 1996). These pupae all die during early to mid-pupal stages or as pharate adults and no flies eclose (Figure 1A). The pupal lethality and head eversion defects observed in AdarG-overexpressing progeny indicate a disruption in ecdysone signaling, either through excessive or insufficient pathway activity (Thummel 1996). We also crossed the *Act*5C^ts^*/CyO* line to a *attP40 UAS-Adar^EA^* or *attP40 UAS-Adar^S^* inserts. The phenotypes of *Act5C^ts^> Adar^EA^* progeny were similar to what was observed with AdarG overexpression; pre-pupal lethality and escaper pupae with the head eversion defect (Figure 1B); *Act5C^ts^> Adar^S^* progeny all died before the pupal stage and no pupae were obtained. Evidently, the novel head eversion defect is due to an additional editing-independent effect of Adar G protein overexpression, since it is mimicked by overexpression of catalytically-inactive Adar EA protein We expected that Adar RNA editing activity is increased above normal by AdarG overexpression without this being the cause of lethality, whereas AdarS overexpreesion lethality is probably still due to increased editing activity.

**Figure 1.**
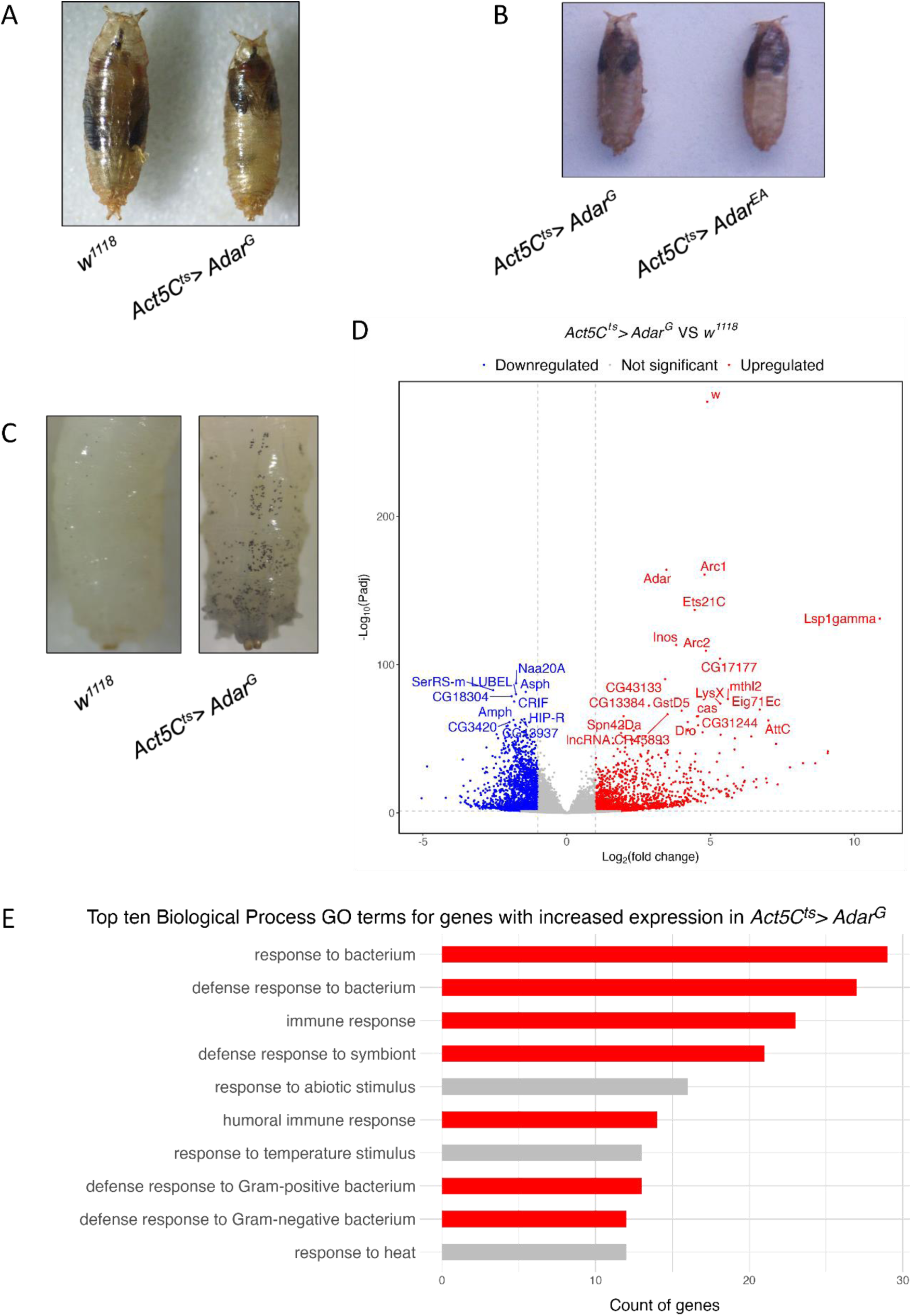
Ubiquitous expression of Adar^G^ throughout development is lethal and leads to aberrant activation of innate immune genes. ***(A)*** The *Act5C^ts^> Adar^G^* pupal lethal phenotype showing truncated legs, smaller wings, extended abdomen region and pupal head eversion defect. ***(B)*** *Act5C^ts^> Adar^EA^* pupae show the same characteristic pupal head eversion defect as AdarG overexpression. ***(C)*** Melanotic spots formed in *Act5C^ts^> Adar^G^* larvae heated to 65 °C for 10 minutes indicate aberrant innate immune induction and aberrant blood cell proliferation with increased melanizing crystal cells. ***(D)*** Volcano plot (-log_10_ (adjusted p value) vs. log_2_(fold change)) for *Act5C^ts^> Adar^G^* vs wild type *w^1118^* pupae RNAseq. ***(E)*** The top 10 enriched Biological Process GO terms for genes that are significantly overexpressed in *Act5C^ts^> Adar^G^* pupae vs wild type pupae. Bars are colored red for immune-related terms.

As a preliminary to a genetic screen for suppressors of the AdarG overexpression defects, *Act5C^ts^/CyO> Adar^G^* virgin females were crossed at 25°C to males of two VDRC RNAi lines homozygous for *UAS-Adar RNAi* transgene insertions. The *Act5C^ts^> Adar*^G^, *Adar RNAi* progeny of this cross, in which the *Act5C^ts^-GAL4* driver directs simultaneous expression of both the *UAS-Adar^G^* and the *UAS-Adar RNAi* constructs, showed nearly complete rescue of the AdarG overexpression lethality (98%) with either VDRC line (Table 1). On the other hand, crosses between *Act5C^ts^/CyO> Adar^G^* virgin females and control males (*w^1118^* or *UAS-GFP*) resulted in 100% lethality of the non-*Cy*, *Act5C^ts^> Adar^G^* progeny. Therefore, the *Adar RNAi* rescue is not due either to reduction in *UAS-Adar^G^* transgene copy number from two in the mothers to one in the progeny since crosses to *w^1118^* did not rescue. It is also not due to weaker GAL4-driven expression of *UAS-Adar^G^* due to the presence of the additional *UAS* in the *Adar RNAi* transgenes, since progeny carrying the *UAS-GFP* transgene do not show rescue.

**Table 1.** Genetic screen for suppressors of the the *Act5C^ts^> Adar^G^* lethal phenotype. *Act5C^ts^/CyO> Adar^G^* virgin females were crossed to *UAS−GFP*, *UAS−Adar RNAi* lines or other RNAi knockdown rescue candidates at 25^°^C.

| Genotype | Rescue (%) |
| --- | --- |
| *Act5C<sup>ts</sup>* > *Adar<sup>G</sup>*, *GFP<sup>14</sup>* | 0.0 |
| *Act5C<sup>ts</sup>* > *Adar<sup>G</sup>*, *Adar RNAi<sup>GD1355</sup>* | 98.3 |
| *Act5C<sup>ts</sup>* > *Adar<sup>G</sup>*, *Adar RNAi<sup>KK100520</sup>* | 98.2 |
| *Act5C<sup>ts</sup>* > *Adar<sup>G</sup>*, *EcRA RNAi<sup>91</sup>* | 69.15 |
| *Act5C<sup>ts</sup>* > *Adar<sup>G</sup>*, *Su(var)205<sup>ZH-21F</sup>* | 46.01 |
| *Act5C<sup>ts</sup>* > *Adar<sup>G</sup>*, *Cam RNAi<sup>KK109037</sup>* | 26.9 |
| *Act5C<sup>ts</sup>* > *Adar<sup>G</sup>*, *Pp2B-14D RNAi<sup>KK107714</sup>* | 17.6 |
| *Act5C<sup>ts</sup>* > *Adar<sup>G</sup>*, *dnc* | 10.3 |
| *Act5C<sup>ts</sup>* > *Adar<sup>G</sup>*, *Smn RNAi<sup>FL26B</sup>* | 8.4 |
| *Act5C<sup>ts</sup>* > *Adar<sup>G</sup>*, *Toll-7 RNAi<sup>GD14417</sup>* | 5.5 |
| *Act5C<sup>ts</sup>* > *Adar<sup>G</sup>*, *Toll-9 RNAi<sup>GD14431</sup>* | 5.2 |
| *Act5C<sup>ts</sup>* > *Adar<sup>G</sup>*, *PPO2 RNAi<sup>KK112850</sup>* | 5.1 |
| *Act5C<sup>ts</sup>* > *Adar<sup>G</sup>*, *Nmdmc RNAi<sup>GD1256</sup>* | 2.9 |
| *Act5C<sup>ts</sup>* > *Adar<sup>G</sup>*, *tub RNAi<sup>VSH330551</sup>* | 2.5 |
| *Act5C<sup>ts</sup>* > *Adar<sup>G</sup>*, *Toll-6 RNAi<sup>GD14438</sup>* | 1.0 |
| *Act5C<sup>ts</sup>* > *Adar<sup>G</sup>*, *Arc1 RNAi<sup>GD16710</sup>* | 0.7 |
| *Act5C<sup>ts</sup>* > *Adar<sup>G</sup>*, *Arc2 RNAi<sup>GD1727</sup>* | 0.0 |
| *Act5C<sup>ts</sup>* > *Adar<sup>G</sup>*, *EcRB1 RNAi<sup>168</sup>* | 0.0 |
| *Act5C<sup>ts</sup>* > *Adar<sup>G</sup>*, *p53 RNAi<sup>GD11134</sup>* | 0.0 |
| *Act5C<sup>ts</sup>* > *Adar<sup>G</sup>*, *Sp7 RNAi<sup>KK111500</sup>* | 0.0 |
| *Act5C<sup>ts</sup>* > *Adar<sup>G</sup>*, *MP1 RNAi<sup>KK107262</sup>* | 0.0 |
| *Act5C<sup>ts</sup>* > *Adar<sup>G</sup>*, *PPO3 RNAi<sup>GD16572</sup>* | 0.0 |
| *Act5C<sup>ts</sup>* > *Adar<sup>G</sup>*, *Myd88 RNAi<sup>GD9716</sup>* | 0.0 |
| *Act5C<sup>ts</sup>* > *Adar<sup>G</sup>*, *pll RNAi<sup>GD1286</sup>* | 0.0 |
| *Act5C<sup>ts</sup>* > *Adar<sup>G</sup>*; *Dif RNAi<sup>KK106594</sup>* | 0.0 |
| *Act5C<sup>ts</sup>* > *Adar<sup>G</sup>*, *Pirk RNAi<sup>KK105790</sup>* | 0.0 |

Given the head eversion defect which is similar in AdarG overexpressing and ecdysone mutant pupae, a genetic screen was conducted, focusing primarily on ecdysone signaling and its upstream regulators: the prothoracicotropic hormone (PTTH) released from brain neurons onto PG cells, calcium signaling downstream of the PTTH/Torso receptor and insulin/TOR growth-regulating pathways (Gilbert et al. 1997; Caldwell et al. 2005; Colombani et al. 2005; Mirth 2005; Niwa and Niwa 2016), (Table 1). Since *Adar^5G1^* null mutants show aberrant innate immune gene induction (Deng et al. 2020), locomotion defects, aberrantly increased TOR activity, defective autophagy and age-dependent neurodegeneration (Khan et al. 2020), genes affecting these pathways were also tested as potential suppressors. Additional RNAi knockdowns and mutants targeting known Adar-edited transcripts expressed in the CNS, particularly those related to calcium signaling and ion channel activity, were also tested (Table 1).

Supporting the involvement of ecdysone signaling in AdarG overexpression lethality, knockdown of the ecdysone receptor A (*EcRA)* isoform in progeny from a cross to an *UAS-EcRA RNAi* line gave 69.15% rescue of AdarG overexpression lethality. Among the different EcR isoforms (Schubiger et al. 2003), only *EcRA* knockdown provided strong rescue, while *EcRB1* knockdown failed to rescue. Knockdown of *calmodulin* (26.9% rescue) or the calmodulin-dependent *Protein phosphatase 2B* (*Pp2B14D*) (17.6% recue) also partially suppressed lethality. Overexpression of *dunce (dnc)*, a cAMP-specific phosphodiesterase, gave 10.3% rescue (Table 1). These results suggest that AdarG overexpression lethality involves aberrant calcium/cAMP signaling. Also consistent with the role of EcR function, knockdown of *survival motor neuron (Smn)* (8.4 % rescue) resulted in partial rescue (Table 1). Smn is an important component of small nuclear ribonucleoproteins (snRNPs) that are essential for splicing. Smn interacts with Gemin proteins (Shpargel et al. 2009; Cauchi et al. 2010), including Gemin5, which is known to associate with nuclear receptors such as EcR, USP and βFTZ-F1, potentially acting as a cofactor in controlling the expression of ecdysone-regulated genes and fine-tuning spatial and temporal responses to ecdysone during development (Gates et al. 2004). These findings support a role for Smn in regulating ecdysone signaling and suggest that AdarG overexpression lethality may be mediated by initially aberrantly enhanced ecdysone pathway activity.

### Aberrantly increased ecdysone signaling and aberrant innate immune induction in AdarG-overexpressing pupa are mediated by EcRA

To characterize the underlying molecular abberations in AdarG overexpressing pupae, whole-transcriptome RNA-Seq was performed on pupae exhibiting the head eversion phenotype. We had noticed that wandering larvae from the *Adar^ts^/CyO> Adar^G^* strain or larvae from crosses of *Adar^ts^/CyO> Adar^G^* to *w^1118^* at 25 °C showed black melanization spots associated with aberrant innate immune induction and associated aberrant proliferation of blood crystal cells. Heating larvae to 65 °C for 10 minutes better visualizes the black spots formed after release of prophenoloxidases 1 and 2 from crystal cells (Figure 1C), which is normally triggered for melanization of parasites.

Differential gene expression analysis on the RNA-Seq data from AdarG overexpressing versus w*^1118^* pupae (Excel Table 1) represented as a Volcano plot of differentially expressed genes (DEGs), show increased expression of Ecdysone genes (*Eig71C*) and innate immune genes *Attacin C* (*AttC*) and *Drosocin* (*Dro*) (Figure 1D). Gene enrichment analysis performed on the 294 most significantly upregulated transcripts (log_2_(fold change) > 3) in AdarG-overexpressing pupae also showed GO term enrichment of immune response pathways (Figure 1E and Supplemental Table S1). Aberrant innate immune induction of antimicrobial peptide (AMP) transcripts occurs in *Adar^5G1^* null mutants but was not previously observed after Adar isoform overexpression (Deng et al. 2020); the immune induction now observed may be another effect of the new higher Adar isoform expression. Surprisingly, ecdysone-related transcripts were not observed in the top ten altered groups in the GO term analysis of the AdarG overexpressing pupa transcriptome. *Eig71* transcripts are elevated in the Volcano plot of differentially expressed genes; these genes encode ecdysone-induced, pupa-specific inate immune defense proteins and also show high induction ratios like AMP transcripts. Evidently, the pupal head eversion defect are associated with smaller expression changes in some ecdysone signaling transcripts.

To validate the transcriptome findings, RT-qPCR was performed on RNA extracted from AdarG overexpressing pupae displaying the head eversion defect or from wildtype *w^1118^* pupae. *Adar* transcript levels are elevated, as expected, in AdarG overexpressing pupae compared to *w^1118^*pupae (Figure 2A), and aberrantly increased expression of ecdysone-related and innate immune AMP transcripts in AdarG overexpressing pupae were confirmed (Fig. 2B-D). We confirmed significant upregulation of transcripts encoding early components of the ecdysone signaling pathway, such as *EcRA* (1.37-fold), *Br-C* (6.22-fold), *E75* (3.44-fold) (Figure 2B) as well as several ecdysone-induced genes, including *Eig71Ec* (268.37-fold), *Eig71Ed* (205.79-fold), *Eig71Eg* (119.58-fold), and *Eig71Eh* (29.43-fold) (Figure 2C). All of the upregulated transcripts, which are normally induced during the pupal stage, encode proteins acting as positive regulators of ecdysone signaling at various hierarchical levels to ensure proper progression of metamorphosis (Ou and King-Jones 2013). In contrast, transcripts encoding mid-to late-stage components of the pathway such as *Hr4* and *ftz-f1* did not show increased expression in AdarG-overexpressing pupae (Supplemental Figure S1). The expression level of FTZ-F1 protein is reduced by aberrantly high ecdysone early in metamorphosis and low FTZ-F1 subsequently leads to lower expression of other late ecdysone signaling proteins. Therefore, the FTZ-F1 level is a regulatory pivot around which aberrantly high expression of early ecdysone signaling proteins can lead to aberrantly low expression of late ecdysone signaling proteins. We also measured the expression of transcripts aberrantly elevated in AdarG-overexpressing pupae in *Act5C^ts^> Adar^G^, Adar RNAi* pupae in which the head eversion defect is rescued and observed restoration of these transcripts to near-normal levels (not shown), although only half of these pupae have the overexpressing genotype and the other half are indistinguishable *CyO* pupae.

**Figure 2.**
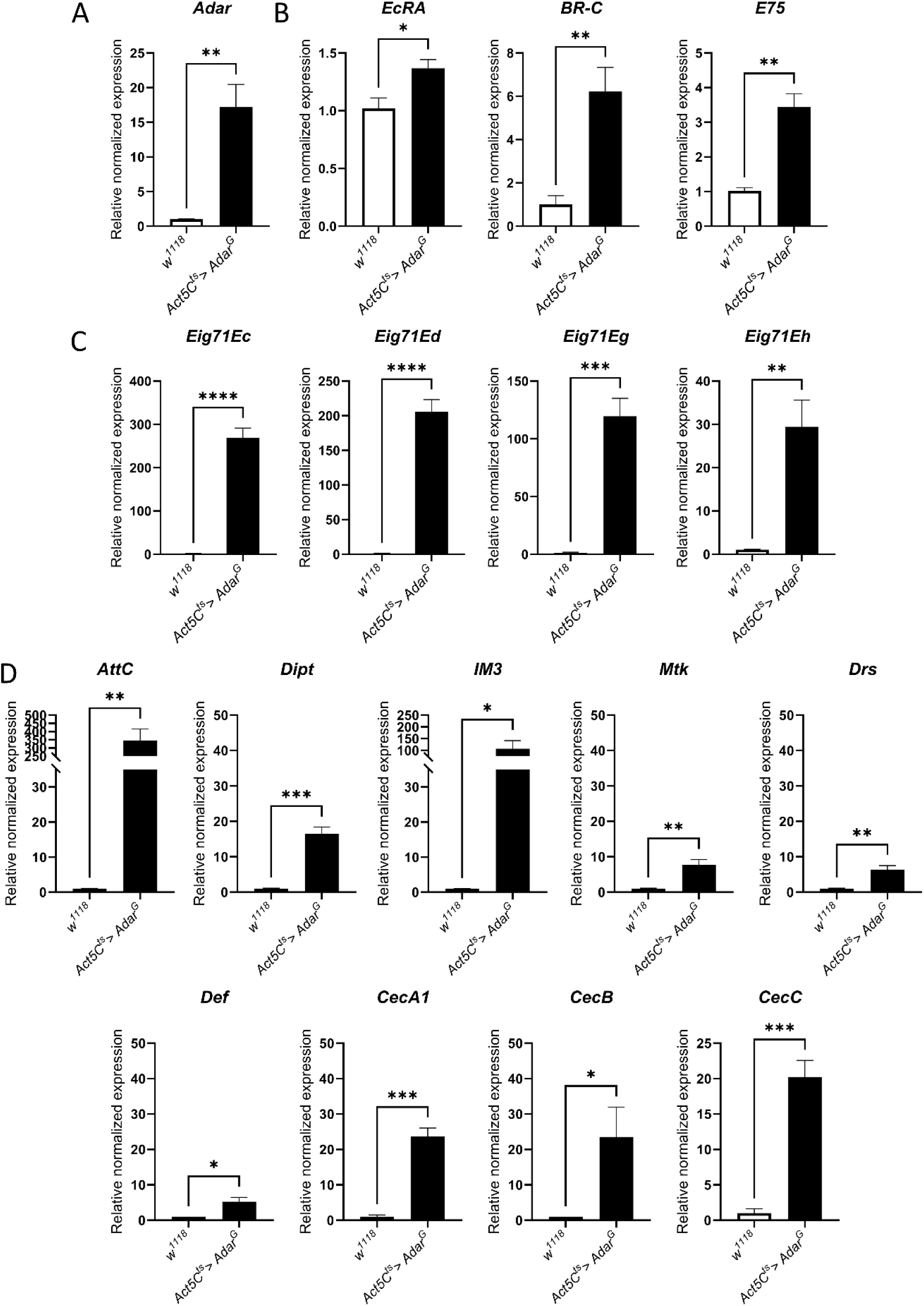
Transcripts encoding ecdysone-related and innate immune proteins are aberrantly elevated in AdarG overexpressing pupae. RT-qPCR on RNA from pupae. Normalized transcript expression levels for the *Act5C^ts^> Adar^G^* relative to wild type. ***(A)*** *Adar* transcript expression level normalized to wild type. ***(B)*** *EcRA* and early ecdysone-response gene (*BR-C* and *E75*) transcript expression levels normalized to wild type. ***(C)*** *Ecdysone-induced gene 71 (Eig71)* transcript expression levels normalized to wild type. ***(D)*** Innate immune AMP transcript expression levels normalized to wild type. The *y*-axis is discontinuous for *AttC* and *IM3*; the break in the *y*-axis represents the jump in the *y*-scale. Below the break, the *y*-scale is the same as that of the other transcripts. *p*-values were calculated by Student’s *t*-test with Welch correction. *: *p*-value < 0.05. **: *p*-value < 0.01. ***: *p*-value < 0.001. ****: *p*-value < 0.0001. ns: not significant. Error bars: SEM (Standard Error of Mean for biological replicates). See Supplemental Table S3 for primers used.

The strong upregulation of AMP transcripts in AdarG-overexpressing pupae detected by RNA-Seq was also confirmed by RT-qPCR. Upregulated transcripts included IMD pathway-regulated transcripts (*Attacin C*, 346.11-fold; *Diptericin*, 16.56-fold), as well as Toll pathway-regulated transcripts (*Drosomycin,* 6.31-fold; *Metchnikowi*n, 7.69-fold; *IM3*, 107.10-fold). Also aberrantly elevated were AMP transcripts regulated by both pathways (*Defensin*, 5.24-fold), and AMPs active against Gram-negative bacteria (*Cecropin A1*, 23.75-fold; *Cecropin B*, 23.54-fold; *Cecropin C*, 20.20-fold), (Figure 2D). Because of the aberrant AMP inductions, the genetic screen for suppressors of AdarG-overexpression effects was extended to incude knocking down key components of the IMD, Toll and JAK-STAT signaling pathways. Surprisingly, knockdown of *Nmdnc/Rel*, *Toll-6*, *Toll-7*, *Toll-9*, *Myd88*, *tube, pelle, Dif* or *Pirk* resulted in little to no rescue of AdarG overexpression lethality nor of the head eversion defect (Table 1). Similarly, knockdown of melanization-related immune effectors**—**including *Prophenoloxidase 3 (PPO3)*, *Serine protease 7 (Sp7)* or *Melanization Protease 1 (MP1)***—**did not give any detectable rescue, while *Prophenoloxidase 2 (PPO2)* knockdown led to only a minimal rescue effect (Table 1). Perhaps it is the elevated ecdysone signaling that drives the aberrant AMP induction or, if the aberrant innate immune induction is the primary defect, then no individual knockdown of an AMP regulator is sufficient to prevent all AMP induction. Transcripts of *Activity-regulated cytoskeleton associated protein 1 (Arc1)* and *2 (Arc2)* were also0 highly upregulated in the RNA-Seq data from AdarG-overexpressing pupae (Figure 1D). Arc1 and Arc2 proteins play roles in mediating intercellular RNA transfer (Ashley et al. 2018; Erlendsson et al. 2020). However, knockdown of *Arc1* and *Arc2* resulted in little or no rescue (Table 1).

We wished to examine effects of rescues in *Act5C^ts^> Adar^G^*, *EcRA RNAi* pupae; however, these rescued pupae are not distinguishable from sibling *CyO* pupae lacking AdarG overexpression. We could distinguish these genotypes in young rescued flies and RT-PCR analyses showed that in *Act5C^ts^ >Adar^G^, EcRA RNAi* compared to *w^1118^* flies the *Adar* transcript is still aberrantly high, as expected (Figure 3A), while *EcRA*, *BR-C* and *E75* are restored to near wildtype levels (Figure 3B). Ecdysone-related transcripts are expected to be lower after pupation but we did not have any AdarG-overexpressing escaper flies to compare to. However, the innate immune AMP transcripts were also restored to near-normal or to below normal levels in *Act5C^ts^ >Adar^G^, EcRA RNAi* compared to *w^1118^* flies (Figure 3C). Ecdysone signaling and innate immune induction interact potently; ecdysone signaling is required to prepare blood cells for innate immune induction after infections and activation of ecdysone receptors during metamorphosis leads to increased expression of antimicrobial peptides (AMPs) (Keith 2023). On the other hand, infection or tissue damage in larvae, leading to aberrant innate immune induction, may delay pupation (Hackney et al. 2012), so as to avoid death during metamorphosis.

**Figure 3.**
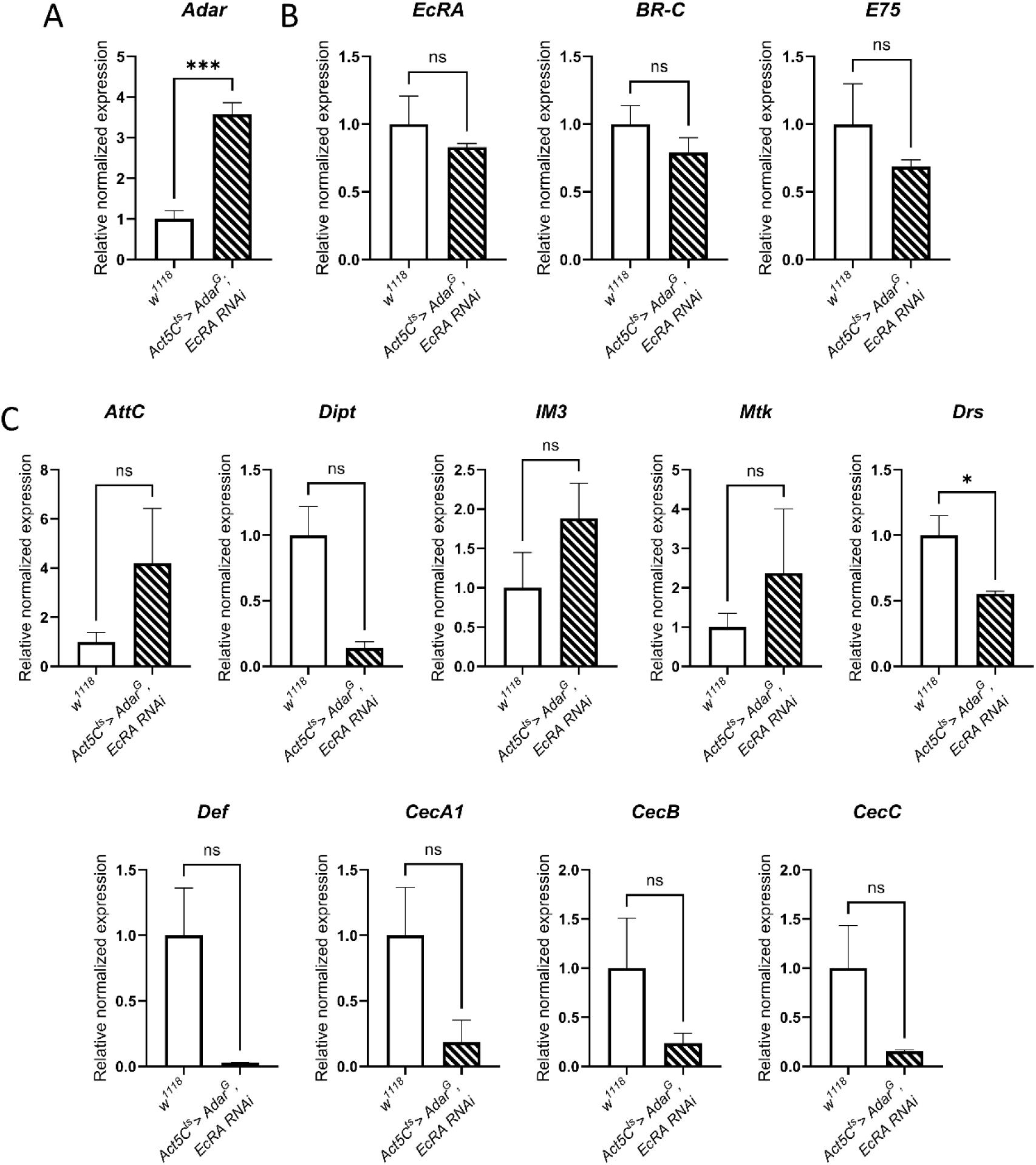
Expression of ecdysone-related and innate immune transcripts is rescued in *Act5C^ts^> Adar^G^, EcRA RNAi* rescued flies. RT-qPCR on RNA from adults. Transcript expression levels in *Act5C^ts^> Adar^G^, EcRA RNAi* relative to wild type. ***(A)*** *Adar* transcript expression level normalized to wild type. ***(B)*** *EcRA* and early ecdysone-response gene (*BR-C* and *E75*) transcript expression levels normalized to wild type. ***(C)*** Innate immune AMP transcript expression levels normalized to wild type transcripts. *p*-values were calculated by Student’s *t*-test with Welch correction. *: *p*-value < 0.05. **: *p*-value < 0.01. ***: *p*-value < 0.001. ns: not significant.Error bars: SEM (Standard Error of Mean for biological replicates). See Supplemental Table S3 for primers used.

We also examined RNA editing at known target sites in the RNA-seq of AdarG-overexpressing versus wildtype *w^1118^* pupae and detected increased editing at a number of sites (Supplementary Excel Table 2 and Supplementary Excel Table 3), which we confirmed by RT-PCR and Sanger sequencing. In *Act5C^ts^> Adar^G^* pupae, editing sites with increased A-to-G editing percentages relative to wild-type were identified. A total of 13 sites across 7 transcripts exhibited an increase in editing of ≥ 30%, with a maximum increase of 40% (Supplemental Table S2, Supplemental Figure S2 and Supplemental Figure S3). All of these sites were also edited in wild-type pupae; we did not detect *de novo* editing sites in AdarG-overexpressing pupae. The AdarG isoform was previously reported to have some slightly different preferences than AdarS for specific editing sites (Jepson et al. 2011); however, such sites are already well edited in *w^1118^*. The increased editing we detected in AdarG overexpressing pupae is mostly at sites with much lower editing in *w^1118^*. Since *Act5C^ts^> Adar^E374A^* mimics the prepupal lethal and pupal head eversion defect of *Act5C^ts^> Adar^G^* the increased editing is not critical for the lethality.

### Adar protein is normally expressed in the prothoracic gland and localizes to pericentric heterochromatin

Due to the ecdysone-related head eversion defect caused by ubiquitous AdarG overexpression, we investigated the expression of Adar in the wildtype prothoracic gland (PG) which produces ecdysone. Although *Adar* is highly expressed in the nervous system, it was previously reported that Adar localizes in neuronal, but not glial nuclei in the larval and adult CNS (Jepson et al. 2011). However, Adar expression was also detected in other tissues, albeit at lower levels (Brown et al. 2014; Leader et al. 2018), including in muscles of *Adar-sGFP* transgenic flies (Sarov et al. 2016; Kanca et al. 2017). We performed immunofluorescent staining and detected Adar-HA protein in the ring gland of *Adar-HA* strain wandering third instar larvae (Figure 4A) with positive staining in the prothoracic gland (PG, outlined in red). In contrast to the neurons, where the expression of Adar is much higher and staining fills the nuclei, in the PG cell nuclei we were able to see a distinct Adar-HA chromosomal localization pattern. When *Adar-HA* larval ring glands were co-stained for Adar-HA and the heterochromatin marker HP1, Adar-HA localized mostly in the HP1-positive areas of pericentric heterochromatin (Figure 4B, white arrows), and was also observed on polytene chromosomes outside the pericentric heterochromatin (yellow arrows). Adar localization to heterochromatic chromosome IV was previously detected in salivary glands of larvae overexpressing Adar-HA (Savva et al. 2013). We also confirmed our findings in *w^1118^, iso31* (Celera wt strain) ring glands stained with antibody against *Drosophila* Adar (Figure 4B, lower panel).

**Figure 4.**
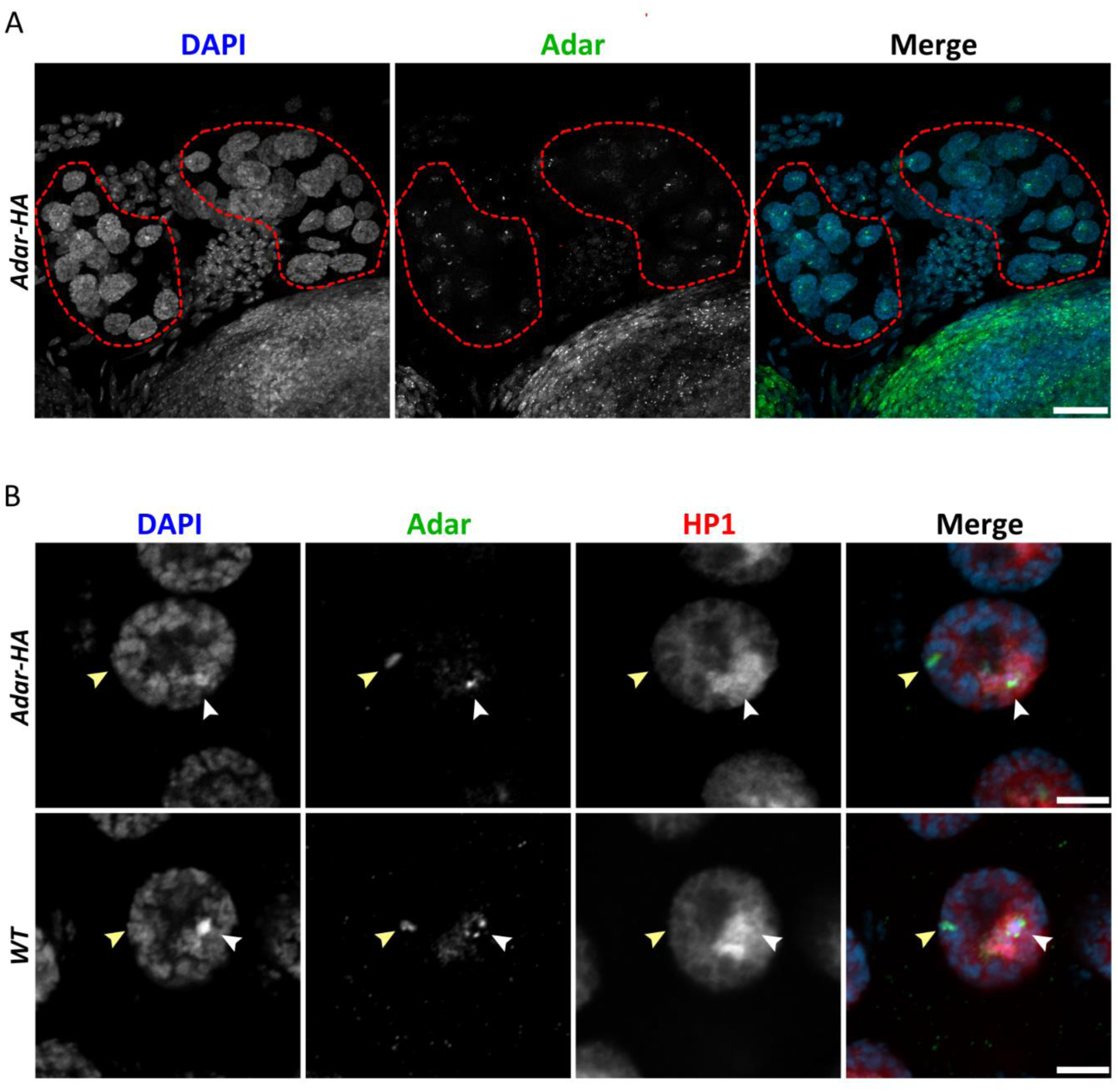
Adar is expressed in the wildtype ring gland and localizes in nuclei, mainly to pericentric heterochromatin. Number of ring glands imaged (n). ***(A)*** Orthogonal projection of the ring gland (n = 3) of *Adar-HA* wandering third instar larvae with the prothoracic gland outlined in red. Adar-HA (green) was visualized using mouse antibody against the HA-epitope tag. DAPI was used to stain DNA (blue). Scale bar, 20 µm. ***(B)*** Confocal sections of prothoracic gland nuclei of *Adar-HA* (n = 4) and *w^1118^ iso31 (WT,* n= 3) third instar wandering larvae stained for Adar (green) and heterochromatin marker HP1 (red). In the *Adar-HA* strain Adar-HA was visualized with a rabbit antibody against the HA-epitope tag. In the case of *WT*, Adar was detected with an antibody against *Drosophila* Adar. Scale bars, 5 µm.

We reanalysed publicly available transcriptome data (GSE76304) on dissected ring glands of Celera *Drosophila* strains (Christesen et al. 2017) and observed *Adar* expression there. To identify Adar RNA editing sites in ring gland transcripts we identified A-to-G changes in cDNAs in the Celera strain which is the source of the *Drosophila* reference genome sequence and free of confounding genomic SNP differences. We found editing at sites already identified in larval transcriptomes, particularly in the *spok* transcript, which is involved in ecdysone biosynthesis and is enriched in PG (Supplementary Excel Table 4).

### AdarG overexpression specifically in the prothoracic gland leads to partial lethality and giant larvae with impaired ecdysone biosynthesis

Increased ecdysone signaling in the AdarG overexpressing pupae suggested possible interactions between Adar and the ecdysone pathway in the prothoracic glands (PG). To explore how Adar regulates ecdysone levels during development we overexpressed AdarG in the prothoracic gland with the Chromosome III *phm22*-*GAL4* driver (Colombani et al. 2005; Mirth 2005); *phantom* (*phm*) encodes an ecdysone biosynthesis enzyme that is very higly expressed in ecdysone-synthesising cells. The *phantom* transcript is three fold more abundant than the *Act5C* transcript in the ring gland transcriptomes (Christesen et al. 2017), even though *phantom* is expressed only in the PG subset of ring gland cells. Surprisingly, *phm22-GAL4*-driven overexpression of the *attP40 UAS-Adar^G^* insertion at 25 °C resulted in partial lethality. The larvae fed for extended periods of time and grew in size and gave rise to bigger pupae and ultimately to bigger surviving flies. However, some animals exhibited lethality in the earlier stages of development, some died during the pupal stages and some died due to failure to eclose (data not shown). A size comparison of *phm22* and *phm22> Adar^G^* pupae is shown in Figure 5A. Again, the phenotypes observed in *phm22> Adar^G^* larvae, pupae and adults were also observed in *phm22> Adar^EA^* animals where Adar EA was expressed from an *attP40 UAS-Adar^EA^* insert (Figure 5A), indicating that the phenotypes are due to overexpression of AdarG or catalytically-inactive AdarEA proteins, and independent of RNA editing activity.

**Figure 5.**
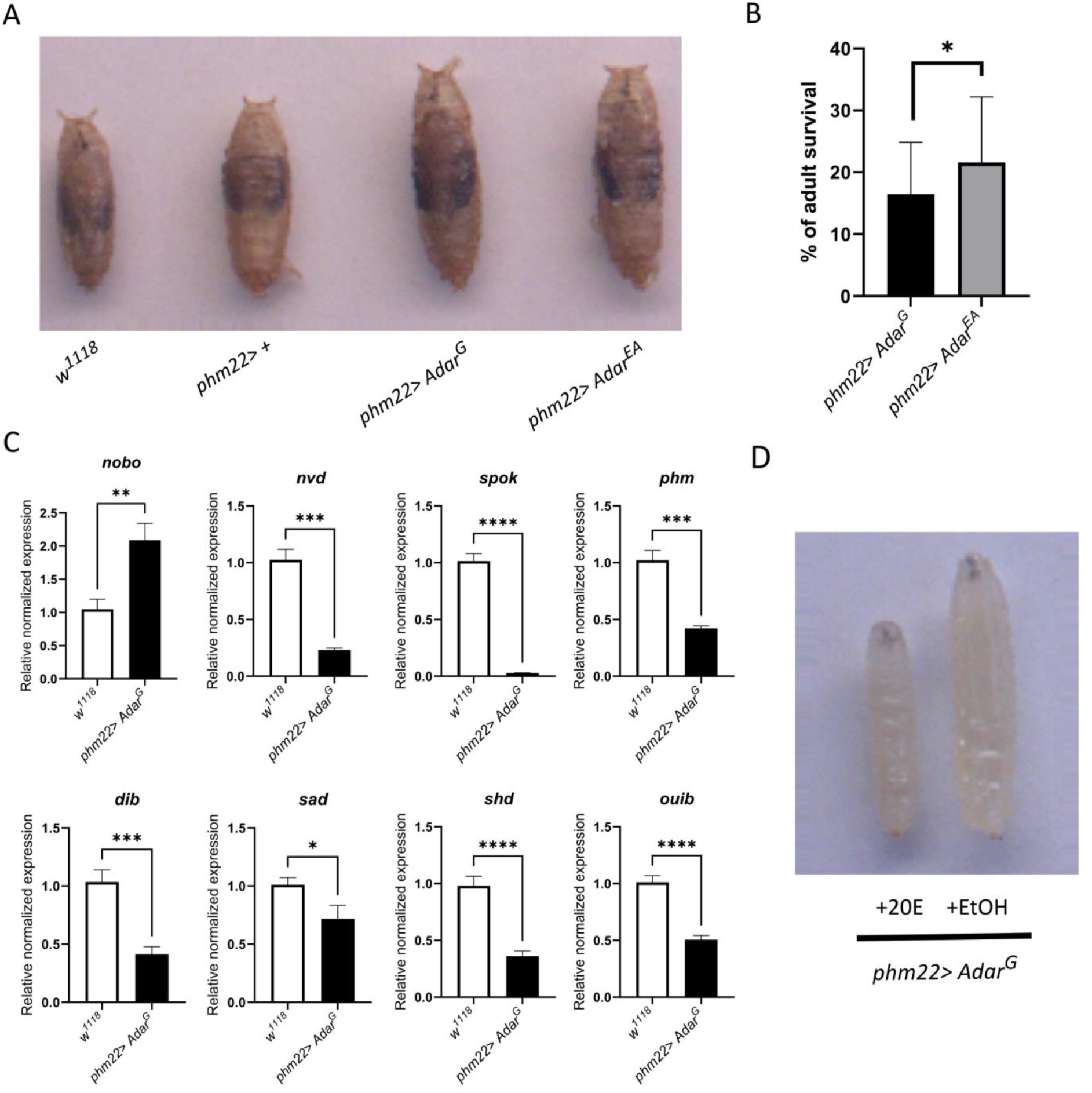
*phm22> Adar^G^* overexpression prevents ecdysone synthesis and gives giant larvae with delayed pupation which are prevented by 20-hydroxyecdysone feeding. ***(A)*** The phenotype of *phm22> Adar^G^* pupae showing giant pupae compared to *w^1118^* and *phm22* pupae. *phm22> Adar^EA^* pupae are similarly giant. ***(B)*** Mean percentage of adult survival of *phm22> Adar^G^* is significantly lower than *phm22> Adar^EA^*. *p*-value was calculated by Fisher’s exact test done on pooled counts of non-balancer and balancer flies. *: *p*-value < 0.05. Error bars: SEM (Standard Error of Mean for biological replicates). ***(C)*** Most transcripts encoding ecdysone biosynthesis enzymes are downregulated in *phm22> Adar^G^* wandering giant larvae. Normalized expression levels relative to wild type for *nobo, nvd, spok, phm, dib, sad, shd* and *ouib* transcripts. *p*-values were calculated by Student’s *t*-test with Welch correction. *: *p*-value < 0.05. **: *p*-value < 0.01. ***: *p*-value < 0.001. ****: *p*-value < 0.0001. ns: not significant. Error bars: SEM (Standard Error of Mean for biological replicates). See Supplemental Table S3 for primers used. ***(D)*** Delayed wandering and development of giant larvae are rescued in *phm22> Adar^G^* third instar larvae when they are grown on food supplemented with 20-hydroxyecdysone (20E, 0,3 mg.g^-1^) versus ethanol (EtOH).

Since both *phm22> Adar^G^* and *phm22> Adar^EA^* showed only partial lethality and similar larval and pupal phenotypes, suggesting that the effect of AdarG is editing-independent, the percentage of AdarG-overexpressing animals eclosing was counted. Virgins of homozygous *phm22-GAL4* were crossed with *UAS-Adar^G^/CyO or UAS-Adar^EA^/CyO* males and survival was measured as the percentage of observed non-*Cy*, AdarG-overexpressing flies compared to their *Cy* siblings. Mean percentages of adult survival of *phm22> Adar^G^* reached 16,47% and of *phm22> Adar^EA^* reached 21,57%. The significance of the difference in survival was addressed by Fisher’s exact test on pooled counts of *CyO* balancer and non-balancer flies between the *Adar^G^* and *Adar^EA^* crosses (Figure 5B). This confirmed a significant difference, suggesting *Adar^EA^* overexpression is slightly less lethal.

The prolonged larval feeding behavior and giant larvae arise from decreased expression of ecdysone biosynthesis genes (McBrayer et al. 2007). Ablation of PTTH neurons inervating PG causes delayed development and size increase as well as lowered expression of ecdysone biosynthesis genes (McBrayer et al. 2007). RNA-Seq analysis (Supplementary Excel Table 5) and RT-qPCR analysis of *phm22> Adar^G^*third instar wandering larvae shows strongly and significantly decreased expression of transcripts encoding ecdysone synthesis enzymes, including *neverland* (*nvd*, 0.23-fold), *spookier* (*spok*, 0.03-fold), *phantom* (*phm*, 0.42-fold), *disembodied* (*dib*, 0.41-fold), *shadow* (*sad*, 0.72-fold), *shade* (*shd*, 0.36-fold) transcripts and the *ouija board* (*ouib*, 0.51-fold) transcript encoding the PG-specific transcription factor for *spok* (Komura-Kawa et al. 2015) (Figure 5C). However, the *Noppera-bo* (*nobo,* 2.09-fold) transcript encoding the glutathione synthetase enzyme required for cholesterol intake into the PG, is upregulated (Enya et al. 2014). Following these findings, we explored the changes in expression of genes encoding the ecdysone signaling proteins that regulate ecdysone production. The level of *EcRA* remained unchanged (Supplemental Figure S4A). However, we observed downregulation of ecdysone early-response gene transcripts including *E75* (0,31-fold), *BR-C* (0,41-fold), *E23* (0,21-fold), *Hr3* (0,35-fold) and *Hr4* (0,09-fold) (Supplemental Figure S4 B, C) as well as of the ecdysone late-response gene *Eig71Ee* (0,37-fold) (Supplemental Figure S4D). The expression of *ftz-f1* (3,53-fold) is upregulated; *ftz-f1* levels are lowered by early pupal ecdysone so increased *ftz-f1* transcript here is consistent with the lowered early ecdysone (Supplemental Figure S4E) (Zhu et al. 2006).

To show that the *phm22> Adar^G^* and *phm22> Adar^EA^* larval developmental arrest phenotype is due to lowered levels of ecdysteroids, larvae were fed food supplemented with 20-hydroxyecdysone (20E, 0,3 mg.g^-1^) or food supplemented with the ethanol (EtOH) diluent. Addition of 20E was able to rescue the *phm22> Adar^G^* prolonged feeding, delay in wandering behavior, and the size increase in third instar wandering larvae (Figure 5D). However, the majority of these larvae were still not able to develop further and died; this is typical for such rescues by ecdysone feeding.

The extreme suppression of ecdysone synthesis in *phm22> Adar^G^* and *phm22> Adar^EA^* are probably due to the very strong *phm22-GAL4* driver. Notably, a specific phenotype was not observed using another PG specific driver, *spok-GAL4*. *Act5C^ts^> Adar^G^* does not cause similar strong inhibition of ecdysone synthesis, so the *phm22> Adar^G^* lethality appears to require the much stronger expression with this driver. The ecdysone signaling pathway has been implicated in muscle remodeling during pupal development (Sandstrom et al. 1997; Zirin et al. 2013; Dequeant et al. 2015) and a group of transient skeletal muscles required for eclosion, the dorsal internal oblique muscles (DIOMs), was reported to be the main source of late pupal ecdysone (Zhang et al. 2024). The *spookier(spok*) transcript is expressed in PG and DIOMs, and is lower than *Act5C* in the ring gland transcriptomes (Christesen et al. 2017).

### Rescue of AdarG overexpression effects by increased expression of *Su(var)205*/HP1 implicates aberrant heterochromatin silencing

The decreased ecdysone production in *phm22> Adar^G^*larvae led us to consider whether Heterochromatin Protein 1 (HP1) might be involved in the aberrant ecdysone-related phenotypes. HP1 mediates heterochromatin silencing but is also required to sustain high expression of the centric-heterochromatin located *spok* and *nvd* genes encoding enzymes required for ecdysone biosynthesis (Ohhara et al. 2022); AdarG expression from an endogenous *Adar* gene modified to express just this isoform has been reported to lower HP1 protein expression and to reduce heterochromatin silencing whereas AdarS increases it (Savva et al. 2013). Therefore, our strong AdarG overexpression might further reduce HP1 and further inhibit HP1-mediated heterochromatin silencing and further decrease the expression of genes such as s*pok* and *nvd*.

We used a Zurich *attP21F UAS-ORF* Chr. II insert to overexpress *Su(var)205*/HP1. Overexpressing HP1 in *phm22> Adar^G^, Su(var)205* did not prevent AdarG ovexpression defects; giant larvae and pupae were still produced. However, expression of *Su(var)205* alone under *phm22-GAL4* driver control in *phm22> Su(var)205* larvae was itself sufficient to cause similar lethality with giant larvae to that seen in *phm22> Adar^G^*. This findings are still consistent with the idea that in *phm22> Adar^G^*, AdarG overexpression decreases HP1, which in turn decreases *spok* expression. Probably, the very high expression levels driven by the *phantom22-GAL4* driver in PG cells is causing overexpressed HP1 to also have a new effect which may be intriguingly similar to that caused by AdarG overexpression.

We also overexpressed HP1 in *Act5C^ts^> Adar^G^, Su(var)205* flies and this did partially rescue the AdarG overexpression lethality (46.01% rescue) (Table 1). We used RT-qPCR to measure expression of *Su(var)205*, *spok* and *nvd* transcripts in AdarG overexpressing pupae (Figure 6A). The *Su(var)205* transcript level is slightly but not significantly increased in AdarG overexpressing pupae, although HP1 protein levels might be more altered. Expression of *spok* transcript is significantly decreased in AdarG overexpressing pupae, consistent with a possible negative effect of AdarG on HP1 function; *nvd* transcript is slightly lower also but not significantly reduced (Figure 6A). In rescued *Act5C^ts^> Adar^G^, Su(var)205* flies the *Su(var)205* transcript is increased 2.45 fold compared to *w^1118^,* as expected (Figure 6B). *Adar* transcript levels remained significantly upregulated in *Act5C^ts^> Adar^G^, Su(var)205* rescued flies, so rescues are not due to reduced levels of AdarG overexpression (Figure 6C). In *Act5C^ts^> Adar^G^, Su(var)205* rescued flies *EcRA* remained upregulated but *BR-C* and *E75* levels were no longer significantly elevated compared to *w^1118^* (Figure 6D), consistent with a possible effect of increased HP1 on ecdysone signaling. Strikingly, AMP transcript levels in *Act5C^ts^> Adar^G^, Su(var)205* surviving adults were returned close to baseline *w^1118^* control levels (Figure 6E). This result shows a very strong impact of HP1 levels on aberrant innate immune induction.

**Figure 6.**
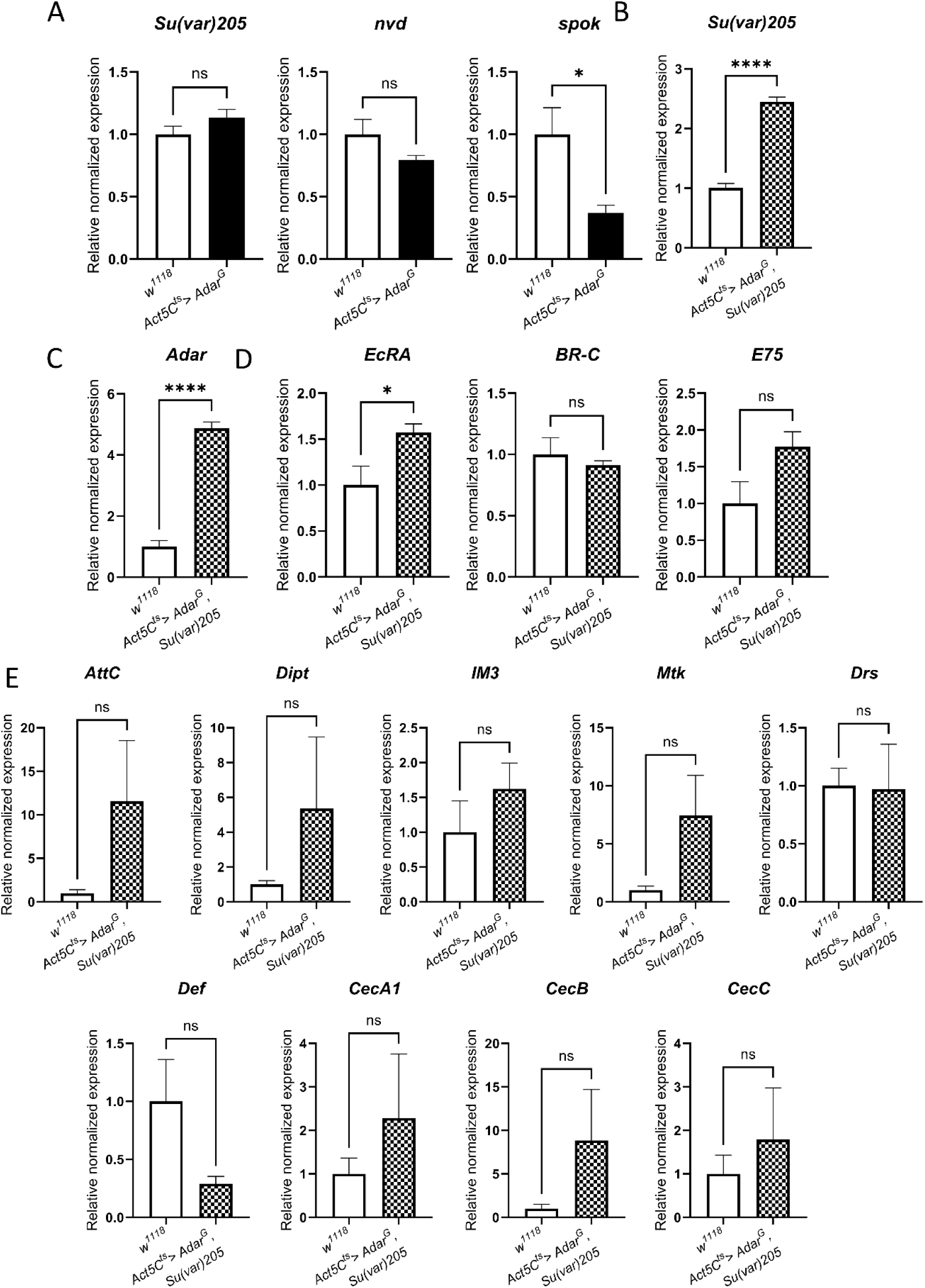
Expression of ecdysone-related and innate immune transcripts are rescued in HP1-overexpressing *Act5C^ts^> Adar^G^, Su(var)205* flies. ***(A)*** RT-qPCR on RNA from *Act5C^ts^> Adar^G^*and wild type, *w^1118^* pupae. Expression levels of *Su(var)205, nvd* and *spok* centric heterochromatin transcripts normalized to wild type. ***(B-E)*** RT-qPCR on RNA from young rescued *Act5C^ts^> Adar^G^, Su(var)205* and wild type *w^1118^* flies. ***(B)*** *Su(var)205* transcript expression level normalized to wild type. ***(C)*** *Adar* transcript expression level normalized to wild type. ***(D)*** *EcRA* and early ecdysone-response gene (*BR-C* and *E75*) transcript expression levels normalized to wild type. ***(E)*** Innate immune AMP transcript expression levels normalized to wild type. *p*-values were calculated by Student’s *t*-test with Welch correction. *: *p*-value < 0.05. **: *p*-value < 0.01. ***: *p*-value < 0.001. ****: *p*-value < 0.0001. ns: not significant. Error bars: SEM (Standard Error of Mean for biological replicates). See Supplemental Table S3 for primers used.

## Discussion

We overexpressed active or inactive Adar proteins at high levels throughout development. Adar proteins are normally expressed at lower levels in embryos and larvae, increase robustly in pupae during metamorphosis and are high in flies, especially in the fly brain. We found that overexpression of AdarG or catalytically inactive AdarE374A proteins under the control of a temperature-regulated *Act5C^ts^-GAL4* driver at 25^°^C gives pre-pupal lethality and pupal head eclosion defects whereas AdarS overexpression gives no pupae. This shows that high level expression of AdarG or AdarEA proteins reveals an editing-independent effect that probably involves Adar interactions with other proteins.

RNA sequencing and qRT-PCR on the RNA extracted from AdarG overexpressing pupae with head eversion defects revealed expected upregulation of transcripts encoding proteins regulating ecdysone synthesis, as well as much stronger upregulation of innate immune AMP gene transcripts. AdarG-overexpressing larvae also have aberrant blood cell proliferation and increased numbers of crystal cells containing melanization enzyme, all indicative of aberrant innate immune induction. The *Adar^5G1^* null mutant also shows aberrant innate immune induction without ecdysone-related defects (Deng et al. 2020), so the new AdarG overexpression pupal head eclosion defect is due to an editing-independent effect of elevated AdarG protein.

We performed a genetic screen for suppressors of AdarG overexpression lethality and of the pupal head eversion defect, particularly using RNAi knockdowns to reduce expression of positive regulators of ecdysone synthesis. Knockdown of the ecdysone receptor EcRA isoform gave the most potent rescue of the *Adar^G^* overexpression lethality, preventing prepupal lethality and the head eversion defect as well as normalizing expression of innate immune and ecdysone-related genes in young rescued flies. It is impossible to state whether the aberrant innate immune induction causes aberrant ecdysone signaling or vice versa. These two signaling pathways interact (Keith 2023), and the two categories of defects arise together in AdarG overexpressing animals.

We show that Adar is normally expressed in nuclei of PG cells that synthesise ecdysone. Adar protein localises to polytene chromomosomes in PG cells, especially to centric heterochromatin. Even though Adar is normally expressed in PGs, overexpression of AdarG or AdarEA under the control of the PG-specific *phantom22-GAL4* driver severely reduced ecdysone synthesis gene expression and gave giant larvae with delayed pupation that were rescued by ecdysone feeding.

We consided whether AdarG overexpression defects could involve aberrant heterochromatin silencing. AdarG was reported to reduce levels of HP1 and to reduce heterochromatin silencing at a cluster of *Hoppel* transposon inserts which generate *Hoppel* dsRNA to initate silencing, wheras AdarS enhances silencing (Savva et al. 2013). Because of this and also because HP1 is required to maintain full expression of the centric heterochromatin-located genes *spookier* and *neverland* encoding key ecdysone biosynthesis enzymes, we examined whether impaired HP1 levels or activity occur in AdarG overexpressing animals. We found that increasing HP1 expression gave a forty six percent rescue of *Act^ts^-GAL4*-driven AdarG overexpression lethality with normalization of ecdysone transcript and innate immune transcript expression.

In the *Adar^5G1^* null mutant the aberrant innate immune induction is mediated through Dicer2 acting as an antiviral dsRNA sensor. Adar protein antagonizes the unknown process by which the Dcr2 antiviral dsRNA sensor leads to transcriptional activation of innate immune genes (Deddouche et al. 2008; Deng et al. 2020). *Drosophila* Adar isoforms also assist or antagonize two types of Dcr2 mediated RNA silencing, cytoplasmic RNA interference (Heale et al. 2009) and nuclear epigenetic transcriptional silencing/ heterochromatin silencing of transposons and some specific genes (Savva et al. 2013). Therefore, the aberrant innate immune induction in AdarG overexpressing animals could result from an excessive AdarG interaction with Dicer2 that somehow also leads to aberrant Dicer2 sensor activation. However, neither *Dcr2 RNAi* knockdown nor any other knockdown of an innate immune regulator, rescued AdarG overexpression lethality. There is evidence of ADAR1/Dicer interaction in vertebrates (Ota et al. 2013) and of Adar-Dcr2 interaction in virus-infected *Drosophila* cells (Rousseau et al. 2025), but further studies will be required to determine the involvenment of Dcr2 in AdarG overexpression lethality, how Adar-Dcr2 interactions occur and how Adar isoforms modulate Dcr2 activity.

HP1 mediates silencing of transposons and host transcripts downstream of dsRNA/Dcr2/Ago2-mediated epigenetic silencing initiation. The AdarG overexpression lethality rescue by increased HP1 is very interesting since the pathway by which the Dcr2 dsRNA sensor activates transcription of the innate immune genes is unknown. There may be a dsRNA/Dcr2/Ago2-mediated regulation of innate immune gene induction which Adar isoforms assist or inhibit. HP1 certainly also affects expression of innate immune genes as increased expression of HP1 in larvae protects against a virus infection by increasing induction of AMPs through a process that remains to be characterized (Wu and Yan 2022).

How do the AdarS and AdarG isoforms exert different effects? *Adar S/G* site editing changes an Adar deaminase domain residue located at the base of a flexible protein loop near the deaminase active site (Figure 7) (Pettersen et al. 2021). In a crystal structure of the ADAR2 deaminase domain monomer the central region of this S/G loop is structured only as far as the residue before the serine (Macbeth et al. 2005). It may be that glycine in AdarG makes the S/G loop more flexible and difficult to fold against the deaminase domain. AdarG editing activity eightfold less than AdarS in vitro at 37 ℃ (Keegan et al. 2005), consistent with an increased entropic energy cost of folding the S/G loop in AdarG. In flies engineered to express either AdarS or AdarG alone, AdarG is only slightly less active at editing sites across the transcriptome than AdarS. This smaller difference in flies at 25 ℃ is consistent with the idea that the difference between AdarS and AdarG is due to to folding of the deaminase domain S/G loop (Keegan et al. 2005; Savva et al. 2012). This temperature-dependent effect would be similar to proposed effects of editing events changing residues to alter protein surface loop flexibilities for temperature adaptation of ion channel responses in Cephalopods (Garrett and Rosenthal 2012).

**Figure 7.**
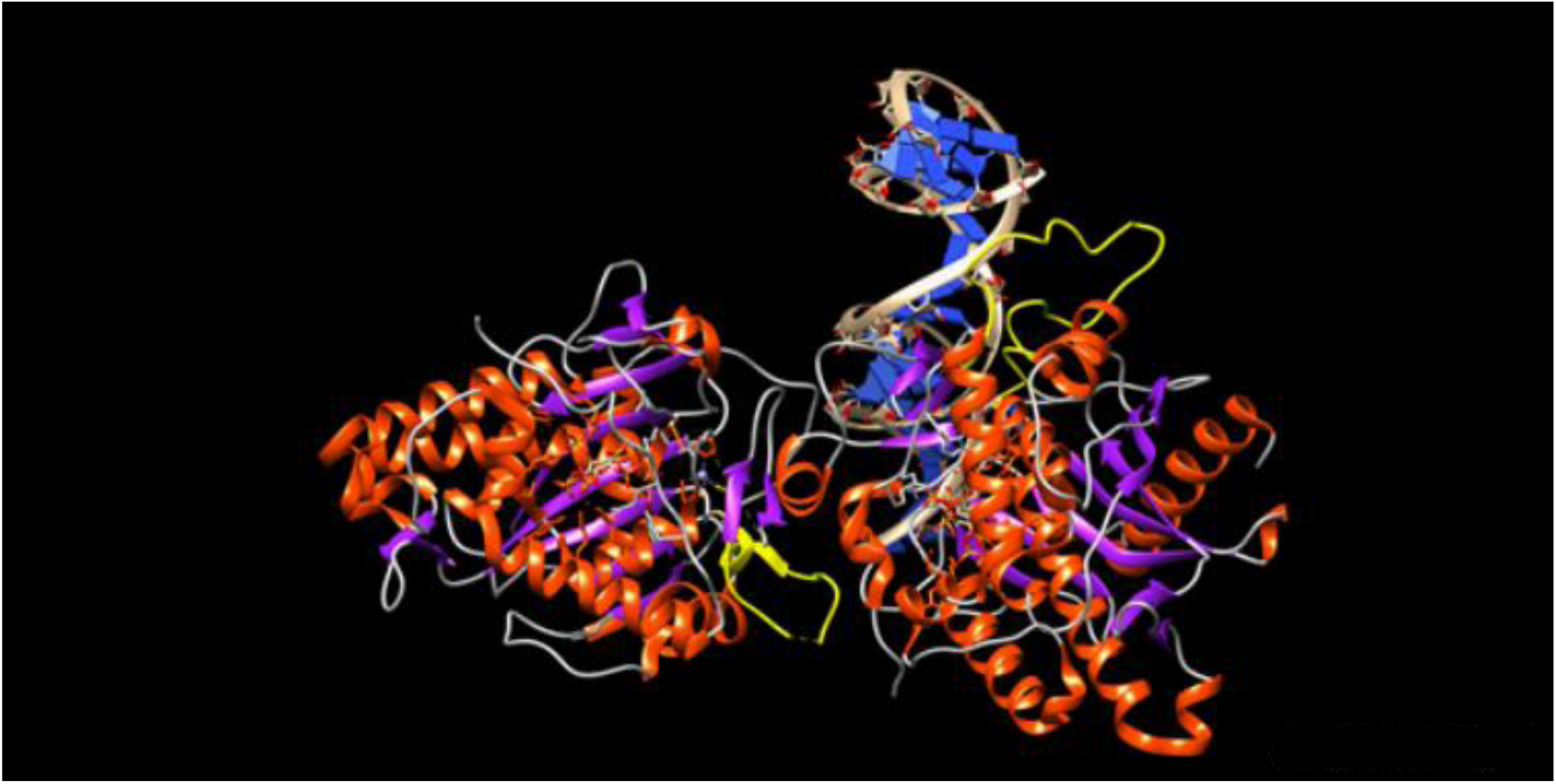
Locations of deaminase domain S/G loops on the cocrystal structure of ADAR2 deaminase domain plus dsRBD2 on dsRNA. The S/G loops coming out from the catalytic site are in yellow, with the backbone at the edited serine residue position in green. The catalytic deaminase domain on the right uses the S/G loop to contact dsRNA. On the other hand, the dsRNA-binding face of the non-catalytic deaminase domain on the left is used for asymmetric dimer formation, which also involves the S/G loop. The non-catalytic deaminase domain S/G loop is not fully folded and the distal part of it is missing from the cocrystal structure. The position of this S/G loop should enable it to contact other proteins on dsRNA.

In a cocrystal of the ADAR2 deaminase domain plus dsRBD2 with dsRNA, the deaminase domains form an asymmetric dimer and the two S/G loops are differently structured for different purposes (Figure 7). In the catalytically active deaminase domain the S/G loop is fully structured and makes dsRNA phosphate contacts. In the non-catalytic deaminase domain the S/G loop is near the asymmetric dimerization site and is structued up to the residue after the serine and but still unstructured further out. The presence of either S or G is likely to affect S/G loop folding in each deaminase domain but the S residue itself does not contact either dsRNA nor the other deaminase domain in the cocrystal structure; these contacts are made by residues further out or futher back along the S/G loop. The S/G loop on the non-catalytic deaminase domain should be available to contact other proteins such as Dicer 2.

## Materials and methods

### *Drosophila* culture and stocks

Fly stocks were maintained at 18 °C on a 12h light/dark cycle and were raised on standard corn meal agar medium. For experimental crosses involving *Act5C-GAL4/CyO* or *Act5C^ts^/CyO>Adar^G^* flies, parental flies were reared on fly food supplemented with organic grape juice diluted 1:1. This food also contained increased nipagin. Crosses were set up on the same grape juice food. For crosses involving *phm22-GAL4* flies, standard food without grape juice and with a normal concentration of nipagin was used.

### Drosophila strains

*w^1118^*, *UAS−Adar^G^ attP2 /TM3, Sb* (this paper), *UAS−Adar^G^ attP40 /SMB9, Cy* (this paper), *UAS−Adar^EA^ attP40 /SMB9, Cy* (this paper), *Act5C−GAL4, Tub−GAL80^ts^(T7) /SM5, Cy; UAS−Adar^G^*(*Act5C^ts^/CyO> Adar^G^*, this paper), *Adar−HA* and *w^1118^; iso31* (Jepson et al. 2011; Robinson et al. 2016) (a kind gift from William J. Joiner, Department of Pharmacology, University of California San Diego), *UAS−GFP^14^*(Bloomington Drosophila Stock Centre, BDSC #4775), *phm22−GAL4* (BDSC #80577), *UAS−Adar RNAi^GD1355^* (Vienna Drosophila Resource Center, VDRC #7764), *UAS−Adar RNAi^KK100520^* (VDRC #105612), *UAS−EcRA RNAi^91^* (BDSC #9328), *UAS−EcRB1 RNAi^168^*(BDSC #9329), *UAS−Smn RNAi^FL26B^* (Chang et al. 2008) (a kind gift from Spyros Artavanis-Tsakonas, Department of Cell Biology, Harvard Medical School), *UAS−Cam RNAi^KK109037^* (VDRC #102004), *UAS−Pp2B-14D RNAi^KK107714^* (VDRC #103144), *UAS−dnc* (Cheung et al. 1999) (a kind gift from Henrike Scholz, University of Cologne), *UAS−Toll-6 RNAi^GD14438^* (VDRC #27102), *UAS−Toll-7 RNAi^GD14417^* (VDRC #39176), *UAS−Toll-9 RNAi^GD14431^* (VDRC #36308), *UAS−PPO2 RNAi^KK112850^* (VDRC #107772), *UAS−PPO3 RNAi^GD16572^*(VDRC #50737), *UAS−Nmdmc RNAi^GD1256^* (VDRC #5706), *UAS−tub RNAi^VSH330551^*(VDRC #330551), *UAS−pll RNAi^GD1286^* (VDRC #2889), *UAS−Myd88 RNAi^GD9716^*(VDRC #25402), *UAS−p53 RNAi^GD11134^* (VDRC #45139), *UAS−Dif RNAi^KK106594^*(VDRC #100537), *UAS−Pirk RNAi^KK105790^* (VDRC #108269), *UAS−MP1 RNAi^KK107262^* (VDRC #104308), *UAS−Sp7 RNAi^KK111500^*(VDRC #104634), *UAS−Arc1 RNAi^GD16710^* (VDRC #48131), *UAS−Arc2 RNAi^GD1727^* (VDRC #3464), *UAS−Su(var)205^ZH-21F^* (Zurich ORFome Project, FlyORF #F003398).

### Generation of *attP* UAS transgenic flies

Sequences encoding *Adar^G^* or *Adar^EA^*were cloned into the *pUASg.attB* plasmid. The transgenic strain *UAS−Adar^G^ attP2/TM3, Sb* was generated by microinjection of pUASg.attB*−Adar^G^* plasmid and PhiC31 integrase-mediated transgenesis (Groth et al. 2004; Fish et al. 2007), resulting in insertion at the *attP2* landing site on chromosome III. To generate the transgenic strains *UAS−Adar^G^ attP40/SMB9, CyO* and *UAS−Adar^EA^/SMB9, CyO,* the pUASg.attB*−Adar^G^* and pUASg.attB*−Adar^EA^*plasmids were sent to the Fly Facility at the Department of Genetics, University of Cambridge, for microinjection and PhiC31 integrase-mediated transgenesis, resulting in insertion at the *attP40* landing site on chromosome II.

### Genetic screen

The *Act5C^ts^/CyO> Adar^G^* recombined stock was generated by combining *Act5C-GAL4*, *TubGAL80^ts^(T7*) on chromosome II with *UAS−Adar^G^* on chromosome III and was subsequently used for the genetic screen. The stock was generated and maintained at 18°C in bottles, with *SM5, CyO* and *TM3, Sb* as balancers for chromosome II and III, respectively. Typically, five virgin females were crossed separately to tester lines at 25 °C, and progeny were screened for the presence of non-*Cy* flies, which indicated either an *Adar RNAi* rescue or a suppressor of *Adar* over expression lethality. The rescue percentage was calculated as the ratio of observed rescue flies to the expected rescue flies, based on the number of progeny carrying the *CyO* balancer chromosome. When the *UAS-GeneX RNAi* tester strain is homozygous for the test construct then the expected number of progeny for a complete rescue is the same as the number of sibling *Cy* progeny from the cross. If the tester construct is heterozygous over some balancer then the expected maximum possible number of rescued flies is half the number of the sibling *Cy* progeny.

### Adult survival

Virgin females of *phm22-GAL4* were crossed to males carrying *UAS-Adar^G^ attP40/CyO* or *UAS-Adar^EA^ attP40/CyO*. Flies were allowed to mate for 5 days, after which the numbers of balancer and non-balancer progeny were scored. The survival percentage of adult flies overexpressing *Adar* isoforms in the prothoracic gland was counted as the ratio of the number of non-balancer progeny to *CyO* balancer progeny. Mean survival percentage ± SEM was plotted. Statistical significance of survival differences between genotypes was assessed by comparing pooled non-balancer and balancer progeny counts with Fisher’s exact test.

### Gene expression analysis

Three samples of total RNA was extracted from 20 pupae each of the desired genotype with the RNAeasy Mini Kit (QIAGEN, #74104) and treated with TURBO DNase (Thermo Fisher Scientific, #AM2238) according to the manufacturer’s instructions. The samples were then sent to Novogene for directional mRNA library preparation with poly(A) enrichment and Illumina paired-end RNA sequencing. Diffrential gene expression was performed as following: raw sequencing reads were quality-checked with FastQC, pre-processed with Cutadapt (v4.3), and aligned to the *Drosophila melanogaster* reference genome (BDGP6.46) with the STAR aligner (v2.7.10a) with Ensembl release 111 gene annotation. Gene counts were estimated with RSEM, and differential expression (DE) analysis was conducted in R with DESeq2. Differentially expressed genes were identified based on an adjusted p-value of < 0.05 and a log2(fold change) threshold of ± 1.

### GO term analysis

Gene Ontology (GO) term enrichment analysis was performed with the enrichGO function from the clusterProfiler R package. Genes with a log₂ fold change greater than 3 in the *w^1118^; Act5C^ts^> Adar^G^* vs wild type comparison were selected for analysis. The org.Dm.eg.db annotation package was used as the reference database, with gene symbols as the key type and the Biological Process (BP) ontology considered. P-values were adjusted for multiple hypothesis testing with the Benjamini-Hochberg method, and only significantly enriched GO terms (adjusted p-value < 0.05) were retained. The results were visualized in bar plots (Figure 1E), where immune-related GO terms were highlighted in red.

### qPCR

For larvae and adult flies, samples of total RNA were extracted from 5-10 larvae or flies each of the desired genotype with TRI Reagent (Sigma-Aldrich, #T9424) and treated with with TURBO DNase (Thermo Fisher Scientific, #AM2238) followed by phenol-chloroform precipitation, according to the manufacturer’s instructions. For pupae, the same total RNA prepared for gene expression analysis was used. cDNA was synthesized with the RevertAid Reverse Transcriptase kit (Thermo Fisher Scientific, #0442) with Oligo(dT) primers. qPCR reactions were performed with the FastStart Universal Syber Green Master (Rox) (Roche, #FSUSGMMRO) and the primers listes in Supplemental Table S3 on a LightCycler 480 II System (Roche). All primers were tested to ensure optimal efficiency. Gene expression levels were normalized to the housekeeping gene *rp49* and referenced to the average *w^1118^* values. Statistical analyses were performed with GraphPad Prism (version 9) with an unpaired t-test incorporating Welch’s correction. A minimum of five and a maximum of nine biological replicates were used, with three technical replicates per gene in each biological replicate.

### Editing analysis

The raw Illumina RNA sequencing data (150 bp paired-end, strand oriented) in fastq format were inspected for quality control during the pre-processing and trimmed for adapters and low-quality sequences (quality score of 20 for at least 70% of the read length was used as a cut-off) with fastp (Chen 2023). The cleaned reads were subsequently aligned against the *Drosophila melanogaster* (dm6) reference genome using STAR (Dobin et al. 2013) with default parameters providing as input the dm6 splice junction list from ncbi RefSeq. To detect the editing sites and quantify the editing level on each of the bam files, the REDItoolDnaRna tool, part of the specific REDItools python tool suite (Lo Giudice et al. 2020), was used with default parameters and providing as input a list of known editing sites in *Drosophila* obtained from the literature. Taking advantage of the experimental design (modulation of Adar expression) the list of editing sites obtained was further refined by removing potential SNPs. Possible single nucleotide polymorphisms (SNPs) recorded in the DGRP database (Mackay et al. 2012; Huang et al. 2014) or the *Drosophila* Genome Nexus database (Lack et al. 2015; Lack et al. 2016) were manually filtered out from the list of edited transcripts. Statistical analyses were performed using GraphPad Prism (version 9) with an unpaired t-test incorporating Welch’s correction.

Editing quantification was validated by direct sequencing of PCR product pools generated from cDNA templates with primers located downstream and upstream of the relevant editing site (Supplemental Table S2). To exclude the presence of SNPs, the corresponding genomic DNA (gDNA) sites were also analyzed. gDNA was extracted from 20 pupae of the desired genotype with phenol-chloroform isolation as described by VDRC (https://shop.vbc.ac.at/media/pdfs/GoodQualityGenomicDNA.pdf). The absence of SNPs was confirmed by direct sequencing of PCR product pools from gDNA templates with primers flanking the relevant site.

The same approach was applied to analyze editing of publicly available ring gland (RG) transcriptomic data (GSE76304). Sample SRR3051642 from the Celera strain was excluded from the analysis, as it displayed a markedly higher number of editing sites compared to samples SRR3051640 and SRR3051641.

### Immunohistochemistry and confocal microscopy

Immunostaining of larval brain and ring glands was performed as previously described (Wu and Luo 2006; Daul et al. 2010; Imura et al. 2017) with some modifications. Briefly, dissections were performed in cold 1x PBS and coarse dissected larvae were fixed with 4% formaldehyde in PBS for 25 min, permealized with 0.1% PBT-X and blocked with blocking solution (5% normal goat serum in 0.1% PBT-X) for 1 hr. The larvae were then incubated with primary antibodies - mouse anti-HA 1:800 (#2367, Cell Signaling), rabbit anti-HA (#37247, Cell Signaling) 1:800, mouse anti-HP1 (#C1A9, DSHB) 1:100, guinea pig anti-dADAR 1:500 (a kind gift from Stephen Goodwin, University of Oxford) overnight at 4°C. After washing with 0.1% PBT-X, tissue was incubated with secondary antibodies and counterstain (Alexa Fluor 488 goat anti-mouse IgG #A11001, Alexa Fluor 568 goat anti-rabbit IgG #A11011, Alexa Fluor goat 488 anti-guinea pig IgG #A11073, Alexa Fluor 647 goat anti-mouse IgG1 #A2124, DAPI #D1306, Invitrogen) diluted to 1:500 and 1:7000 in blocking solution for 1.5 hr. Following washing in 0,1% PBT-X and 1x PBS, the brain and ring gland complexes were dissected in 1x PBS and mounted on poly-L-lysine coated slides in Prolong Glass antifade mountant (#P36980, Invitrogen). Ring glands were imaged with a Zeiss LSM800 confocal microscope with 63x oil objective. Z-stacks were taken with 0.19 µm intervals. Images were processed in ZEN blue and QuickFigures (Mazo 2021).

### 20-hydroxyecdysone feeding

20-hydroxyecdysone (#FH10452, Biosynth) was dissolved in 100% ethanol. Food was supplemented with 20E to final concentration 0.3 mg.g^-1^ (Talamillo et al. 2008). Food with the same volume of ethanol was used as a negative control. Flies were allowed to mate and lay eggs for 5 days. The resulting phenotype of the larvae was scored.

## Supporting information

Supplementary data Hajji et al

Supplementary excel tables Hajji et al

## Acknowledgements

We would like to thank Dr. Goodwin for the Adar antibody. This work was supported by the Czech Science Foundation GAČR 21-27329X (to MO’C), EU Horizon 2020 Research and Innovation Programme, Marie Sklodowska-Curie Grant Agreement No 956810. We acknowledge the core facility CELLIM supported by the Czech-BioImaging large RI project (LM2023050 funded by MEYS CR) for their support with obtaining scientific data presented in this paper. YD was funded by the China Scholarship Council, grant number 202106915017.

## Author Contributions

K.H., D.A, B.N, NS. YD. A.K. and N.S., performed the experiments and analyzed the data L.P.K., and M.A.O. supervised the study. K.H, D.A, B.N, A.K, N.S. M.A.O. and L.P.K. wrote the paper.

