## Supplementary data Hajji et al for "Editing-independent effects of Adar in *Drosophila melanogaster*"

[1]CEITEC Masaryk University, Kamenice 735/5, E35, Brno, 62500, Czech Republic. [2]National Centre for Biomolecular Research, Faculty of Science, Masaryk University, Kamenice 5, 625 00 Brno, Czech Republic. [3]College of Animal Science and Technology, Sichuan Agricultural University, Chengdu 611130, Sichuan, China. [4]Department of Biology, Faculty of Medicine, Masaryk University, Brno, Czech Republic. [5]Department of Biosciences, Biotechnology and Environment, University of Bari "Aldo Moro", Via Orabona 4, Bari, 70124, Italy. [6]Institute of Biomembranes, Bioenergetics and Molecular Biotechnologies, National Research Council of Italy, Via Giovanni Amendola, Bari, 122/O, 70126, Italy

**Supplemental Table S1. List of transcripts included in the top 10 enriched Biological Process GO terms for genes that are significantly overexpressed in *Act5C^ts^> Adar^G^* pupae vs wild type pupae.**

| GO term | Increased transcripts |
|---|---|
| response to bacterium | *PGRP-SB2; AttC; Eig71Ec; Eig71Ed; wntD; Eig71Eh; Eig71Ei; Eig71Ef; Listericin; LysX; Eig71Eg; Ets21C; Dro; AttB; CecB; CecA2; PGRP-SC1b; TotM; LysP; TotZ; DptB; CecC; edin; AttA; hemo; DptA; CecA1; GNBP-like3; Npc2e* |
| defense response to bacterium | *PGRP-SB2; AttC; Eig71Ec; Eig71Ed; wntD; Eig71Eh; Eig71Ei; Eig71Ef; Listericin; LysX; Eig71Eg; Ets21C; Dro; AttB; CecB; CecA2; PGRP-SC1b; TotM; LysP; DptB; CecC; edin; AttA; hemo; DptA; CecA1; Npc2e* |
| immune response | *PGRP-SB2; AttC; CG5246; upd3; CG5909; Dro; AttB; CecB; CG18180; CecA2; PGRP-SC1b; CG14499; CG18179; Eig71Ee; DptB; CecC; CG8329; edin; AttA; CG33465; DptA; CecA1; GNBP-like3* |
| defense response to symbiont | *PGRP-SB2; AttC; CG5246; Listericin; upd3; CG5909; Dro; AttB; CecB; CG18180; CecA2; PGRP-SC1b; CG18179; Eig71Ee; CecC; CG8329; AttA; CG33465; DptA; CecA1; GNBP-like3* |
| response to abiotic stimulus | *Hsp70Ba; Fst; w; Hsp70Aa; TotM; Hsp70Bb; Hsp70Bbb; TotZ; Hsp22; Hsp70Ab; Hsp26; Adar; AttA; Hsp70Bc; DptA; amn* |

| | |
|---|---|
| humoral immune response | *AttC; upd3; Dro; AttB; CecB; CecA2; PGRP-SC1b; Eig71Ee; DptB; CecC; edin; AttA; DptA; CecA1* |
| response to temperature stimulus | *Hsp70Ba; Fst; Hsp70Aa; TotM; Hsp70Bb; Hsp70Bbb; TotZ; Hsp22; Hsp70Ab; Hsp26; Adar; Hsp70Bc; amn* |
| defense response to Gram-positive bacterium | *AttC; wntD; Listericin; Dro; AttB; CecB; CecA2; TotM; DptB; CecC; edin; AttA; CecA1* |
| defense response to Gram-negative bacterium | *Listericin; LysX; Dro; CecB; CecA2; PGRP-SC1b; LysP; CecC; edin; AttA; DptA; CecA1* |
| response to heat | *Hsp70Ba; Hsp70Aa; TotM; Hsp70Bb; Hsp70Bbb; TotZ; Hsp22; Hsp70Ab; Hsp26; Adar; Hsp70Bc; amn* |

**Supplemental Table S2. List of transcripts with more than a thirty percent increase in A-to-I editing in *Act5C*$^{ts}$> *Adar*$^{G}$ pupae compared to wild type.**

| Transcript | Genomic site | Increased editing (%) | Region | Amino acid change and additional notes |
|---|---|---|---|---|
| *pico* | X:19835017 | +32% | exonic | K604R, both residues are positively charged |
| *Mhc* | 2L:16780080 | +34% | intronic | |
| *CAP* | 2R:10280988 | +31% | exonic | S550G, outside of functional domain |
| *Prosap* | 2R:14062507 | +30% | exonic | Q1812R, inside of a polyQ region |
| *dnr1* | 2R:22564377 | +37% | exonic | I592V, outside of functional domain |
| *sls* | 3L:2039908 | +35% | 3’UTR | |
| *sls* | 3L:2039918 | +31% | 3’UTR | |
| *CG12581* | 3R:4185912 | +33% | 3’UTR | |
| *CG12581* | 3R:4185926 | +30% | 3’UTR | |
| *CG12581* | 3R:4185928 | +34% | 3’UTR | |
| *CG12581* | 3R:4186053 | +37% | 3’UTR | |
| *CG12581* | 3R:4186057 | +30% | 3’UTR | |
| *CG12581* | 3R:4186091 | +40% | 3’UTR | |

**Supplemental Table S3. List of primers used in this study.**

| Name | Sequence | Source |
|---|---|---|
| *Eig71Ec* fwd | CTCGGTGCGAATTGTCTCTG | (Zsindely et al. 2009) |
| *Eig71Ec* rev | ACGGGTAGTTGGGGTCCTAC | |
| *Eig71Ed* fwd | ATGTGAACGCTGTGTGGAAA | |
| *Eig71Ed* rev | GCCAGCGAGTTCAGCAATA | |
| *Eig71Eg* fwd | TGGCTTTCTGCTGCATATTG | |
| *Eig71Eg* rev | CCAGCTCACAACGGGTTAAT | |
| *Eig71Eh* fwd | TGACTGTCTGCTTCCTGGTG | |
| *Eig71Eh* rev | CCTGGAGTTTGGAGTCAC | |
| *BR-C* fwd | TAACCTCGGCGTTCGAGAATC | FlyPrimerBank |
| *BR-C* rev | TTTGCAGGGTGTGCTCTTGA | |
| *E75* fwd | ATGCAACAGAGCACCCAGAAT | |
| *E75* rev | CATGGAGTAGGAGGGGCAATC | |
| *Hr4* fwd | GTCGAGCCTCTCCCGATTC | |
| *Hr4* rev | CGTATCACACTCATACCAGCC | |
| *ftz-f1* fwd | CCTACTGCCGATTCCAGAAGT | |
| *ftz-f1* rev | GTCCACCACGCATTCTATCCG | |
| *rp49* fwd | CCGCTTCAAGGGACAGTATC | (Deng et al. 2020) |
| *rp49* rev | GACAATCTCCTTGCGCTTCT | |
| *Adar* fwd | GGCTATAACCGAAAATTGCCACA | |
| *Adar* rev | TGTCTTAGCTCATTCAGCATGG | |
| *AttC* fwd | TTGGGTGGATCACTCACATC | |
| *AttC* rev | GCGTATGGGTTTTGGTCAGT | |

| | | |
|---|---|---|
| *Dipt* fwd | ACCGCAGTACCCACTCAATC | |
| *Dipt* rev | ACTTTCCAGCTCGGTTCTGA | |
| *Def* fwd | GCTATCGCTTTTGCTCTGCT | |
| *Def* rev | GGTGTGGTTCCAGTTCCACT | |
| *IM3* fwd | ACTCGCCTTCGTTTTGGGTC | |
| *IM3* rev | TTAGGCCCTCACATTGCAGA | |
| *Mtk* fwd | TACATCAGTGCTGGCAGAGC | |
| *Mtk* rev | AATAAATTGGACCCGGTCTTG | |
| *Drs* fwd | CTCCGTGAGAACCTTTTCCA | |
| *Drs* rev | ACAGGTCTCGTTGTCCCAGA | |
| *Su(var)205* fwd | AGTACGCCGTGGAAAAGATCA | (Zenk et al. 2021) |
| *Su(var)205* rev | CGTGTTCTCAGTTTCGGGATAG | |
| *EcRA* fwd | GACTTGATGTTACCGAGGACTGGATG | |
| *EcRA* rev | GCAGGTCCCTTCTTGCTCTTCTT | |
| *CecA1* fwd | TCTGGCCATCACCATTGGAC | |
| *CecA1* rev | GACATTGGCGGCTTGTTGAG | |
| *CecB* fwd | CGTCTTTGTGGCACTCATCCT | |
| *CecB* rev | ATTCCGAGGACCTGGATTGA | |
| *CecC* fwd | GATCTTCGTTTTCGTCGCCC | |
| *CecC* rev | GCGCAATTCCCAGTCCTTGA | |
| *CG12581* fwd | CGGAGAAATGGGACTGAAAGC | |
| *CG12581* rev | GTGGAAGAATATTTCAACGGAGCT | |
| *sls* fwd | AAGGCGGATCCAGGACAAAG | |
| *sls* rev | TCCCATTGATAGCGATGCCC | |

| | |
|---|---|
| *Prosap* fwd | ACCTATGGTGGTTCGCCTTG |
| *Prosap* rev | GGCAACAACAATCTGCAGCA |

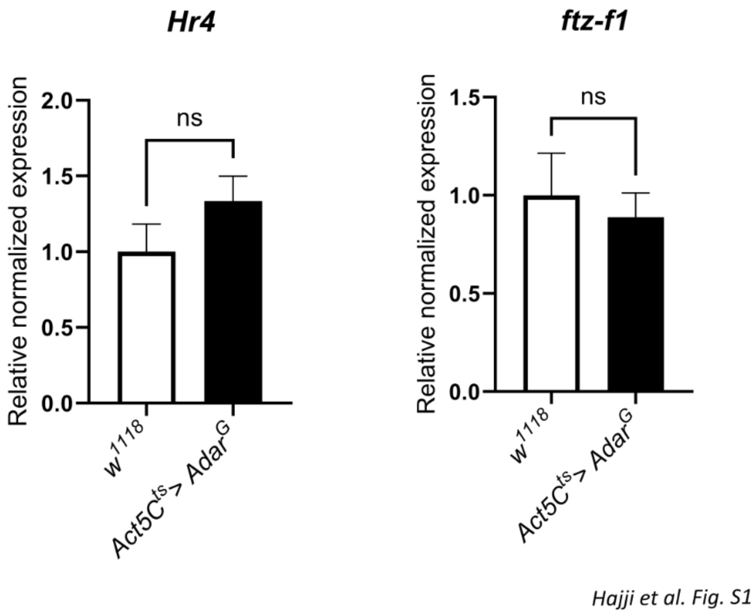


**Supplemental Figure S1. Mid-late ecdysone response gene (*Hr4* and *ftz-f1*) transcripts are not aberrantly elevated in AdarG overexpressing pupae**. RT-qPCR on RNA from pupae. Normalized expression levels of *Hr4* and *ftz-f1* transcripts in $Act5C^{ts}$> $Adar^{G}$ relative to $w^{1118}$ wild type. *p*-values were calculated by Student's *t*-test with Welch correction. ns: not significant. Error bars: SEM (Standard Error of Mean for biological replicates).

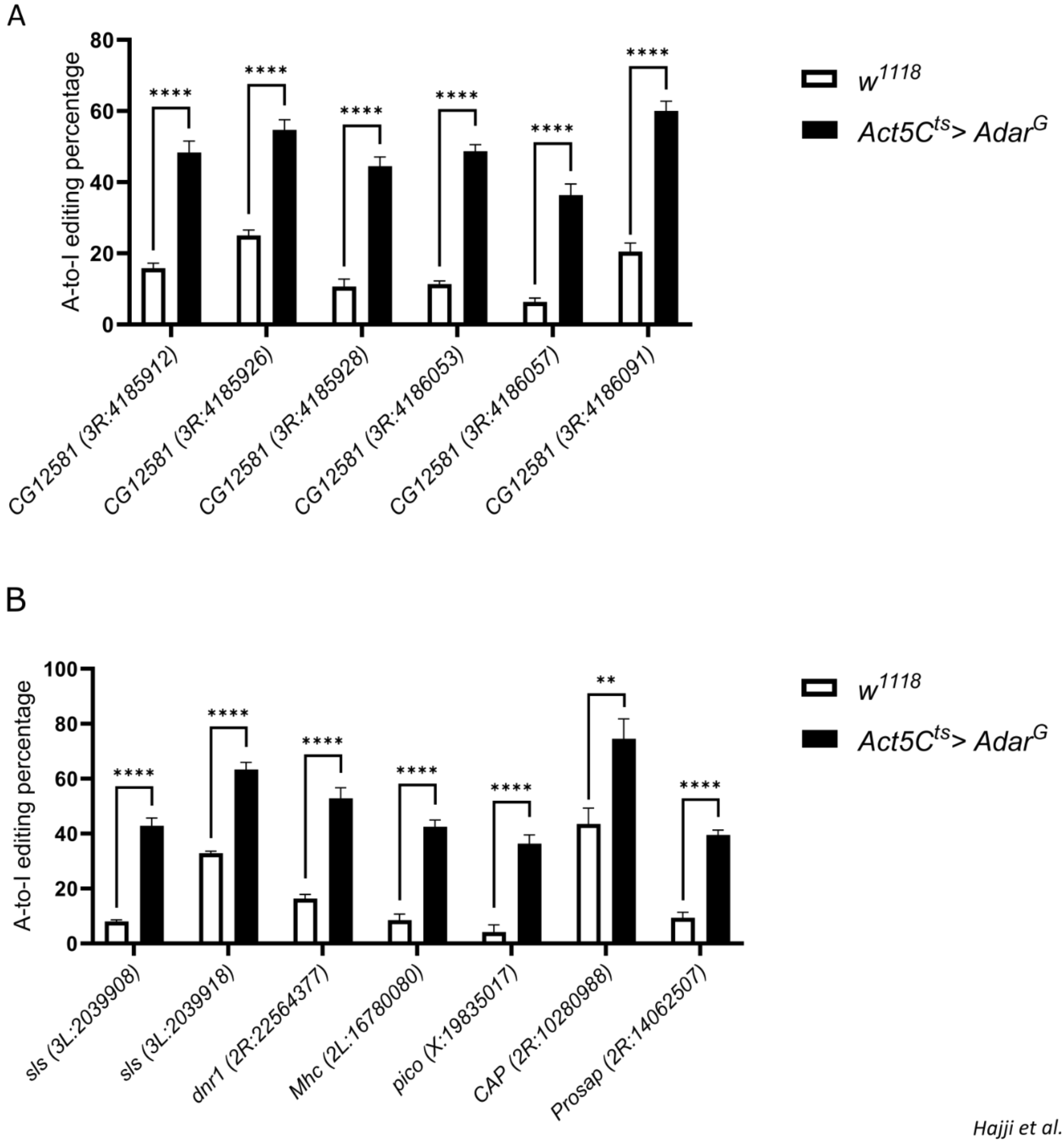


*Hajji et al. Fig. S2*

**Supplemental Figure S2. Sanger sequencing confirmation of increased editing in AdarG overexpressing pupae.** ***(A)*** Editing analysis for *CG12581* transcript. ***(B)*** Editing analysis for *sls*, *dnr1*, *Mhc*, *pico*, *CAP* and *Prosap* transcripts. *p*-values were calculated by Student's *t*-test with Welch correction. *: *p*-value < 0.05. **: *p*-value < 0.01. ***: *p*-value < 0.001. ****: *p*-value < 0.0001. ns: not significant. Error bars: SEM (Standard Error of Mean for biological replicates).

Six of the edting sites were located in the 3'UTR of the uncharacterized transcript *CG12581*. Two additional sites were located in the 3'UTR of *sallimus (sls)*, and one site was found in an intron of *Myosin heavy chain (Mhc)*, both of which encode key structural and contractile muscle proteins. Four editing sites were located in coding regions of four different transcripts: *pico*, *Cbl-associated protein (CAP)*, *Prosap* and *defense repressor 1 (dnr1)*, resulting in single amino acid substitutions. In *pico*, the *Drosophila* ortholog of mammalian lamellipodin (Lyulcheva et al. 2008), the editing-induced substitution K604R was identified, in which lysine at position 604 was replaced by arginine - both positively charged residues. In *CAP*, a cytoskeletal adaptor and vinculin-binding partner (Bharadwaj et al. 2013), an S550G substitution was observed. This residue lies outside of the known SH3 domains and annotated protein regions. Editing in *Prosap*, a scaffolding protein of the SHANK family located in postsynaptic densities (Liebl and Featherstone 2008), resulted in a Q1812R substitution within a polyQ stretch outside of functional domains. Finally, in *dnr1*, an inhibitor of the IMD signaling pathway (Primrose et al. 2007; Guntermann et al. 2009), increased editing led to an I592V substitution outside the protein’s RING-type domain.

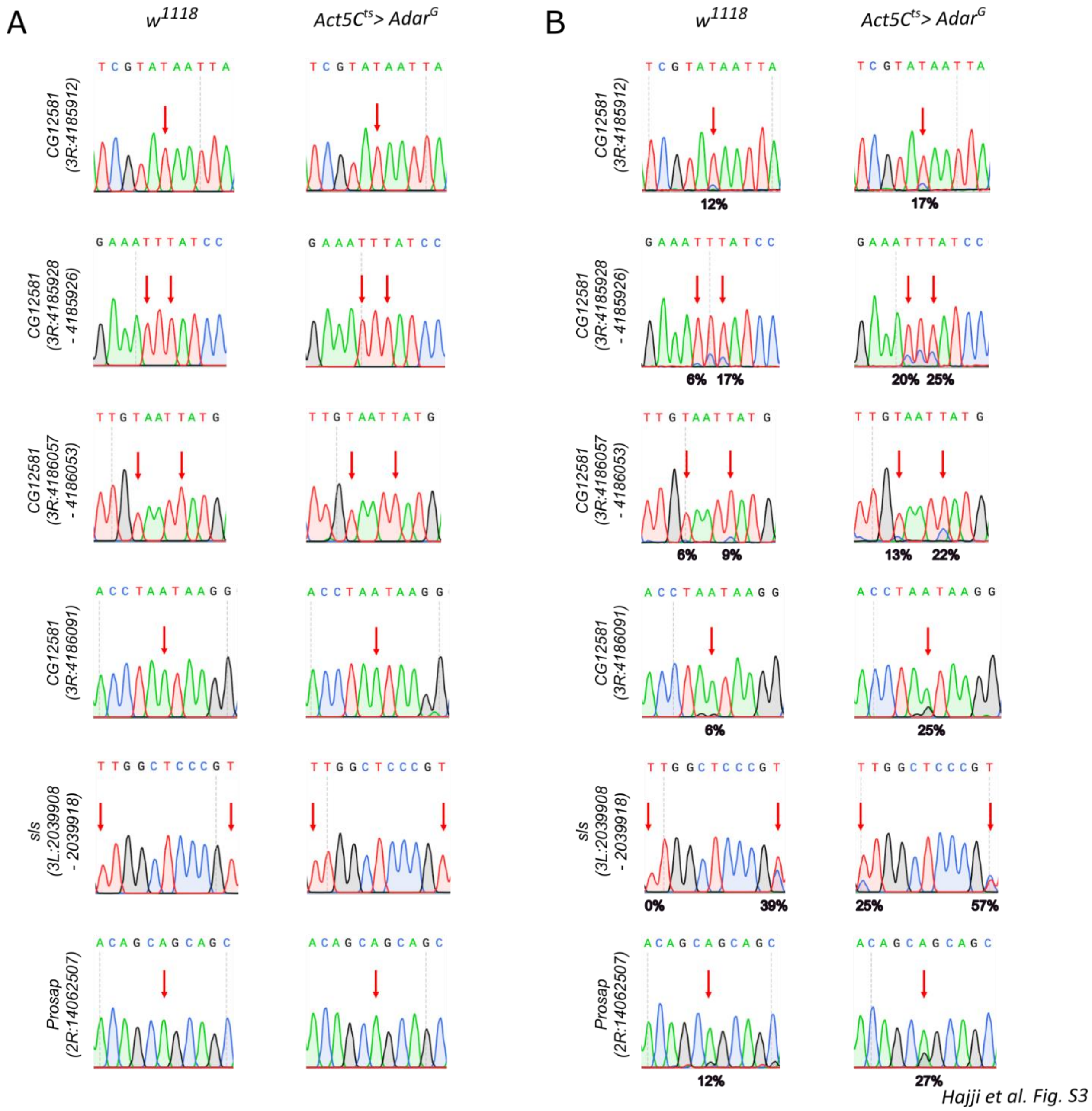


Hajji et al. Fig. S3

**Supplemental Figure S3. Sanger sequencing validation of increased editing in AdarG overexpressing pupae.** ***(A)*** Representative Sanger sequencing chromatograms showing genomic DNA sites corresponding to RNA hyper-edited sites in *Act5C$^{ts}$> Adar$^{G}$* pupae compared to wild type. The site of interest is highlighted with a red arrow. ***(B)*** Representative Sanger sequencing chromatograms showing RNA editing of transcripts that are hyper-edited in *Act5C$^{ts}$> Adar$^{G}$* pupae compared to wild type. The edited site is highlighted with a red arrow.

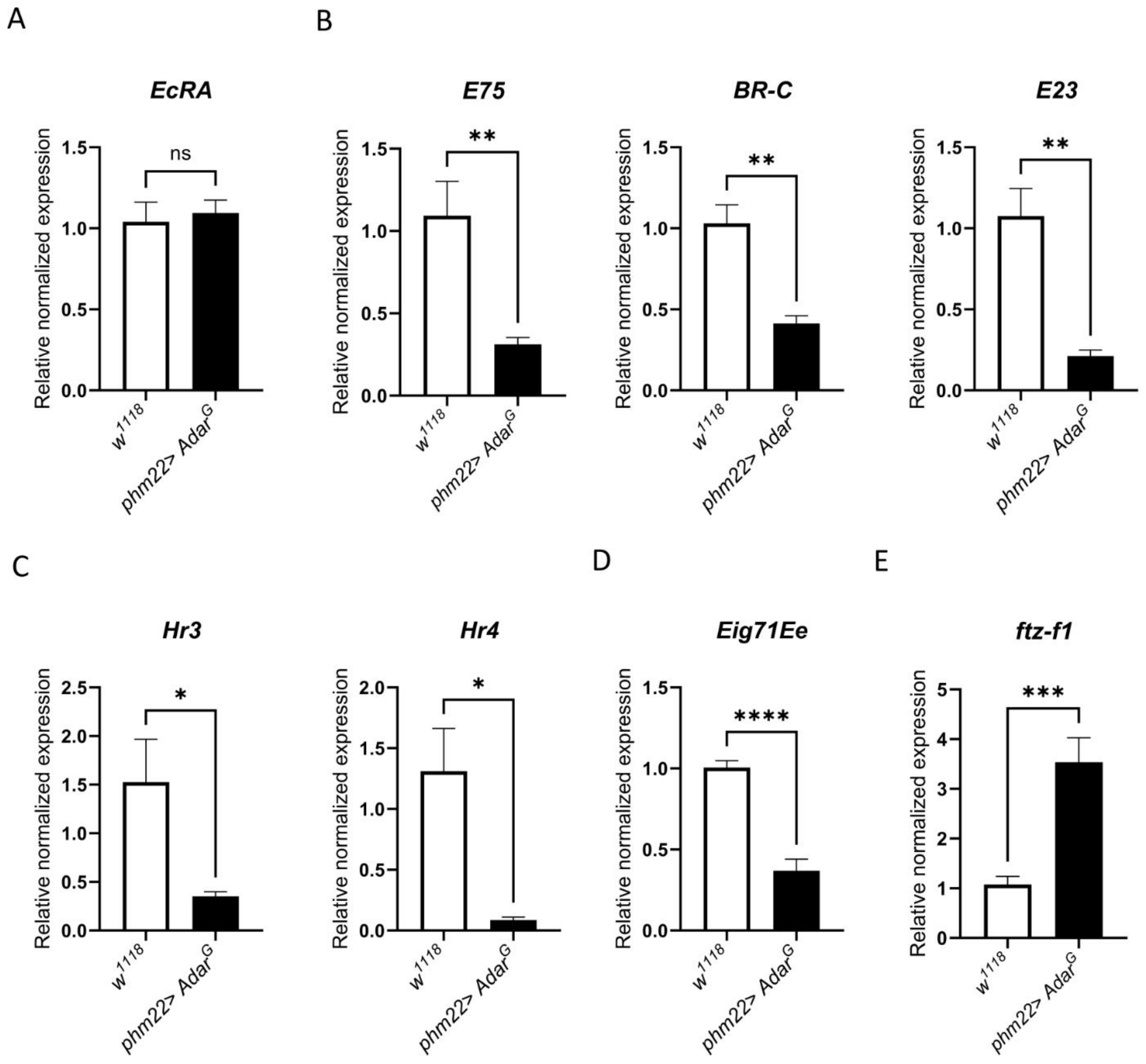


*Hajji et al. Fig. S4*

**Supplemental Figure S4. Levels of transcripts encoding ecdysone signaling proteins measured in RNA from *w*$^{1118}$ and *phm22> Adar*$^{G}$ giant wandering larvae.** ***(A)*** *EcRA* transcript expression level normalized to wild type. ***(B)*** Early ecdysone-response genes (*E75, BR-C, E23*) transcript expression levels normalized to wild type. ***(C)*** Early ecdysone-response gene (*Hr3, Hr4*) transcript expression levels normalized to wild type. ***(D)*** Late ecdysone-response gene *Eig71Ee* transcript expression level normalized to wild type. ***(E)*** *ftz-f1* transcript expression level normalized to wild type. *p*-values were calculated by Student's *t*-test with Welch correction. *: *p*-value < 0.05. **: *p*-value < 0.01. ***: *p*-value < 0.001. ****: *p*-value < 0.0001. ns: not significant. Error bars: SEM (Standard Error of Mean for biological replicates).
